# Intrinsic structural covariation links cerebellum subregions to the cerebral cortex

**DOI:** 10.1101/2024.02.16.580701

**Authors:** Zilong Wang, Jörn Diedrichsen, Karin Saltoun, Christopher Steele, Sheeba Rani Arnold-Anteraper, B.T. Thomas Yeo, Jeremy Schmahmann, Danilo Bzdok

## Abstract

The human cerebellum is increasingly recognized to be involved in non-motor and higher-order cognitive functions. Yet, its ties with the entire cerebral cortex have not been holistically studied in a whole-brain exploration with a unified analytical framework. Here, we characterized dissociable cortical-cerebellar structural covariation patterns across the brain in n=38,527 UK Biobank participants. Our results invigorate previous observations in that important shares of corti-cal-cerebellar structural covariation are described as i) a dissociation between the higher-level cognitive system and lower-level sensorimotor system, as well as ii) an anticorrelation between the visual-attention system and advanced associative networks within the cerebellum. We also discovered a novel pattern of ipsilateral, rather than contralateral, cerebral-cerebellar associations. Furthermore, phenome-wide association assays revealed key phenotypes, including cognitive phenotypes, lifestyle, physical properties, and blood assays, associated with each decomposed covariation pattern, helping to understand their real-world implications. This systems neuroscience view paves the way for future studies to explore the implications of these structural covariations, potentially illuminating new pathways in our understanding of neurological and cognitive disorders.

## Introduction

Traditionally, the cerebellum has been regarded as purely involved in motor control and motor skill acquisition. Yet, accumulating evidence suggested its role in various non-motor as well as higher-order cognitive functions (Leiner et al., 1989; Schmahmann, 1991). Indeed, despite the simple and uniform cellular organization of the cerebellum cortex, a series of tract-tracing studies in monkeys identified multiple closed-loop non-overlapping cortico-ponto/dentato-cerebellar circuits existing in parallel for the prefrontal area 46, posterior parietal area 7b, frontal area 9 and somatomotor cortex (Kelly & Strick, 2003; Middleton & Strick, 1994; Middleton & Strick, 2001; Schmahmann & Pandya, 1989a, 1991a, 1997a). Delineating the cerebrocerebellar polysynaptic connectivity, as done using advanced axon tracer studies in monkeys, is not feasible in the living human brain (Li et al., 2019). Yet, connectivity analyses using functional magnetic resonance imaging (Buckner et al., 2011; Guell, Gabrieli, & Schmahmann, 2018; Ji et al., 2019; King et al., 2023) are consistent with anatomical studies pointing to segregated, partially overlapping loops between different groups of neocortical and cerebellar regions, involving systems supporting both sensorimotor and many non-motor cognitive functions (Schmahmann, 1996; Schmahmann et al., 2019). Here we applied an alternative approach by quantitative modelling of the cortex-cerebellum structural covariation which captures coherent population variation across thousands of people. This approach was naturally sensitive to systematic covariation relationships among multiple cerebrocerebellar regions, which were typically hard to investigate due to their parallel polysynaptic circuit links. This analytical approach traced out the patterns of which cerebellum subregions show structural variation that systematically cooccurs with structural variation in other brain network systems across individuals. To explore real-world relevance, we subsequently delineated the associations between these identified cortex-cerebellum structural covariation patterns (modes) and phenotypic measures indexing behavior, cognition, and physical conditions.

Invasive tract tracing studies provide a unique tool to isolate direct region-region axon pathways. The tract-tracing techniques are designed to meticulously study specific neural pathways in detail (Saleeba et al., 2019). However, this strict localizationist perspective treated the connections from a given brain region at hand as independent from those in other brain regions. Moreover, this technology, by its nature, is not applicable in humans (Li et al., 2019). Furthermore, given the complexity of the brain and limited resources, it is challenging to conduct tract tracing studies that cover larger territories of the brain (Bohland et al., 2009; Osten & Margrie, 2013). In addition, between-species differences are another source of difficulty when extrapolating from cerebellum findings in animals to human biology.

Evolutionarily, as the association cortex, (subserving executive control, language processing and complex social interactions), disproportionately expanded in humans relative to monkeys and apes (Hill et al., 2010; Preuss et al., 2004; Sherwood et al., 2012; Van Essen & Dierker, 2007), parts of the cerebellar cortex (Crus I/II) and cerebellar output dentate nucleus (ventrolateral pro-portion) have similarly enlarged disproportionately (Balsters et al., 2010; Buckner, 2013; Leiner et al., 1986, 1993). Because the cerebral cortex and the cerebellum are highly interconnected, as the cerebral cortex expanded and evolved to become more functionally diverse in humans, it may be expected from the evolutionary perspective that the cerebellum expands and evolves in concert with the cerebral cortex to support those advanced cognitive functions (Whiting & Barton, 2003). Despite the suspected evolutionary co-expansion between specific regions in the cortex and cerebellum, this “big” and “small” cortex of the human brain were routinely studied in isolation (Balsters et al., 2010).

At the level of individual development, the postnatal cortical expansion pattern is known to show certain similarities to the evolutionary trajectory such that expanding higher associative cortical regions, supporting advanced cognitive functions, mature later in life. In contrast, primary visual, auditory and sensorimotor regions appear already mostly mature at birth (Hill et al., 2010). Similarly, cognitive regions of the cerebellum myelinate and mature relatively later (Crus I/II, VIIb, VIIIa) than the cerebellar regions involved in sensorimotor function (Lobule I-IV, VIIIb, IX) (Gaiser et al., 2023; Liu et al., 2022). This is consistent with the observation that the early matured regions support crucial survival functions at an early age, but the formation of advanced cognitive functions is subject to influences through environment and experience throughout life. The cerebellum is postulated to contribute its unique transform to sensorimotor, cognitive and limbic functions (Schmahmann, 1996; Schmahmann et al., 2019), so the coordinated maturation of the cerebellum and the cerebral cortex is crucial for the optimal development of cognitive and motor functions (Kipping et al., 2016). However, previous studies typically investigated only the cerebral cortex (Hill et al., 2010) or only the cerebellar cortex (Liu et al., 2022) in isolation, without considering their interlocking relationships.

The evolutionary and developmental evidence provide support for the prediction that dedicated corresponding region sets across the entire cerebral cortex and cerebellum co-expand and co-mature jointly. If this is correct, it should be possible to observe coherent cerebellum-cortex resonance at the inter-individual level too. In our present investigation, we hypothesized existence of one or more configurations of structural variation that simultaneously occur in a manner that is distributed spatially throughout the cortex-cerebellum complex. Our goal was to holistically chart the covariation between the entire cerebellar system and the entire cortical system at the population level with mission-tailored statistical tools.

Classic anatomical atlases (Angevine et al., 1963; Schmahmann et al., 2000) view the cerebellum on the basis of *structural* landmarks via gross morphological features into the mediolateral division of vermis vs hemispheres, and the anterior lobe (lobules I-V), posterior lobe (lobules VI-IX), and flocculonodular lobe (lobule X). However, recent advances in functional neuroimaging analyses explored segmentation of the cerebellar cortex based on *functional* characteristics, such as patterns of neural activation being similar to other brain networks, or involvement in specific cognitive tasks or processes. In a group-averaged functional connectivity study opting for a winner-takes-all approach (WTA, each voxel is only assigned to a single brain system), Buckner et al. (2011) discovered a roughly homotopic topographical mapping between the cortex and the cerebellum. This correspondence showed that larger higher-order cerebral networks are more extensively represented in the cerebellum. Furthermore, these authors found the majority of the human cerebellar cortex maps to some parts of the cerebral association cortex. The sensorimotor regions accounted for only a small portion of the cerebellum; with the notable exceptions of visual and auditory cortices. Ultimately, the cerebral association networks showed several anterior and posterior correlates in the cerebellum, mirroring the well-established dual representation of the body motor map (Grodd et al., 2001; Habas, Axelrad, & Cabanis, 2004; Habas, Axelrad, et al., 2004; Nitschke et al., 1996; Rijntjes et al., 1999; Schlerf et al., 2010; Snider & Stowell, 1944; Thickbroom et al., 2003). In the WTA approach each cerebellar voxel is assigned to the most correlated brain network, supporting only one possibility for the cortical-cerebellar correspondence. This may have led to an incomplete view of the relationship between cerebral cortex and cerebellum. By embracing a latent-factor approach, we were able to explore the previously untapped possibility that each cerebellar subregion can be linked to several brain phenomena at the same time, and vice versa. Our approach thus allowed for more than one explanation about the role of each subregion in the cerebellum in the context of the entire brain.

To achieve our goals, we mined the UK Biobank (UKBB) population cohort as it provides high-quality structural whole-brain scans for ∼40,000 individuals with ∼1,000 in-depth phenotype measurements. By leveraging an analytical framework for doubly multivariate latent-factor decomposition, we here conjointly investigated the multiple concurrent structural covariation patterns between the entire cerebellar cortex parcellated at 28 region resolution (with a structural and a novel functional atlas) and the entire cerebral cortex (at both the 7-functional-network and 100-subregion granularity) at the population level. We further profiled the key phenotypes associated with each decomposed covariation pattern (mode) to understand their real-world implications by means of phenome-wide association assays.

## Methods

### Population data resource

The UK Biobank (UKBB) is an epidemiology resource that is unusually rich in behavioral and demographic assessments, medical and cognitive measures, as well as biological samples for ∼500,000 participants recruited from across Great Britain (https://www.ukbiobank.ac.uk/). This openly accessible population dataset aims to provide high-quality brain-imaging measurements for ∼100,000 participants. The present study conducted under UK Biobank application number 25163 was based on the recent data release from February/March 2020 that augmented brain-scanning information and expert-curated image-derived phenotypes of gray matter morphology captured by T1-weighted structural MRI of 38,527 participants with 47% men, 53% women and the ages of 40 to 70 years when recruited (54.8±7.5 years). All participants provided written informed consent (for details see http://biobank.ctsu.ox.ac.uk/crystal/field.cgi?id=200).

### Brain imaging and preprocessing procedures

As an attempt to improve comparability and reproducibility, our study built on the uniform data preprocessing pipelines designed and carried out by FMRIB, Oxford University, Oxford, UK (Alfaro-Almagro et al., 2018). Magnetic resonance imaging scanners (3-T Siemens Skyra) were matched at several dedicated data collection sites with the same acquisition protocols and standard Siemens 32-channel radiofrequency receiver head coils. The anonymity of the study participants was protected by defacing the brain scans and removing any identifying information. Automated processing and quality control pipelines were deployed (Alfaro-Almagro et al., 2018; Miller et al., 2016). Noise was filtered out by means of 190 sensitivity features to improve the homogeneity of the imaging data. This approach allowed for the reliable identification and exclusion of brain scans with artefacts, such as excessive head motion.

A three-dimensional (3-D) T1-weighted magnetization-prepared rapid gradient echo (MPRAGE) sequence at 1-mm isotropic resolution was used to obtain structural MRI brain-imaging data as high-resolution images of brain anatomy. Preprocessing included gradient distortion correction (GDC), skull-stripping using the Brain Extraction Tool (Smith, 2002), motion correction using FLIRT (Jenkinson et al., 2002; Jenkinson & Smith, 2001), and nonlinear registration to MNI152 standard space at 1-mm resolution using FNIRT (Andersson et al., 2007). All image transformations were estimated, combined, and applied by a single interpolation step to avoid unnecessary interpolation. Tissue-type segmentation into cerebrospinal fluid (CSF), grey matter (GM), and white matter (WM) was applied using FAST [FMRIB’s Automated Segmentation Tool (Zhang et al., 2001)] to generate full bias-field-corrected images. In turn, SIENAX (Smith et al., 2002) was used to derive volumetric measures normalized for head sizes.

### Target atlas definitions

For the cerebral cortex, GM volume extraction was anatomically guided by the widely used Schaefer-Yeo reference atlas (Schaefer et al., 2018), yielding target features at 100 and 7 parcel resolutions: We extracted 100 cerebral cortical regions across the whole brain for each participant. We next normalized these raw cortical regional GM volumes to represent them as their ratios relative to the total cortical GM volume within each participant. In addition, at the 7-parcel resolution, we took their mean ratio to summarize all the cortical regions belonging to the same predefined canonical network, thus obtaining 7 mean ratios for the 7 networks for each participant. Finally, we z-scored the obtained quantities across participants for both the 100-region variable set and the 7-network variable set.

For the cerebellar cortex, we used two sets of target atlases to derive GM regional volumes for the sake of comparison: a widely used structurally defined atlas (Diedrichsen et al., 2009) based on macro-anatomical landmarks and a recent functionally defined atlas (Zhi et al., 2023) based on neural activity responses across many experimental tasks. This is to evaluate the functional cerebellar parcellation’s effectiveness for explaining the structural covariation across individuals. The first set, sourced directly from the UK Biobank Imaging (UKBB data fields 25893 to 25920), is based on the structurally derived probabilistic SUIT atlas, which uses landmarks such as lobuli and vermis as a point of orientation (Diedrichsen et al., 2009). The structurally derived SUIT atlas divides the entire cerebellum into 28 neighbouring compartments, encompassing the left and right hemispheres of lobules I-IV, V, VI, Crus I, Crus II, VIIb, VIIIa, VIIIb, IX, and X, as well as the vermis of lobules V, VI, Crus I, Crus II, VIIb, VIIIa, VIIIb, IX, and X. The second set’s GM volume extraction was informed by a novel functionally derived probabilistic parcellation of the entire cerebellum, which was based on an aggregate of 7 extensive functional datasets (Nettekoven et al., 2023). For the original version of the atlas, we generated a version with 28 interdigitated functional cerebellar regions (by fusing S2 and S3, as well as A3 and S5 on both the left and right side) to ensure similar number of atlas parcels and thus comparable statistical modeling properties between the two cerebellar atlases. In the end, mirroring the approach with the cortex, we calculated the regional ratios in relation to the total cerebellar GM volume and standardized them across participants for both the lobular anatomical atlas (Diedrichsen et al., 2009) as well as the functional atlas (Nettekoven et al., 2023).

Building upon prior UK Biobank research (Schurz et al., 2021; Spreng et al., 2020), we carried out a preliminary data cleaning step using standard linear regression. In particular, we removed the inter-individual variation in the aforementioned four groups of MRI-derived atlas region measures (i.e., z-scored ratio measures before the regression) that could be explained by the following nuisance variables of no interest: body mass index, head motion during task-related brain scans, head motion during task-unrelated brain scans, head position and receiver coil in the scanner (x, y, and z), position of scanner table, as well as the data acquisition site, in addition to age, age^2^, sex, sex*age, and sex*age^2^. The data cleaning was done by keeping the residuals orthogonal to all the nuisance variables from a linear projection of the z-scored ratio measures to the confounding space (Lindquist et al., 2018). Consequently, the z-scored, denoised regional ratio measures from 100 cortical regions, the mean ratio measures from the 7 Schaefer-Yeo networks (cf. above) and the ratio measures from 28 cerebellar regions - whether derived structurally or functionally - formed the foundation for subsequent analysis steps.

### Population covariation between cerebellar subregions and cortical regions

As the core analysis of our present investigation, we sought to investigate dominant patterns or ‘modes’ of structural covariation that shed light on how structural variation among the cerebellar subregions can explain structural variation among all regions of the cortical brain mantle. Given the need to assess the multivariate relationship between two high-dimensional variable sets, Partial Least Squares Regression (PLSR) was our natural method of reference. A first variable set *X* was constructed from the cerebellar parcels (number of participants × 28 structurally or functionally derived cerebellar parcels matrix). A parallel variable set *Y* was constructed from the cortical subregional gray matter volume ratio or mean network gray matter volume ratio (number of participants × 100 whole-brain cortical parcels matrix or 7 networks matrix).

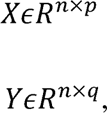

where *n* denotes the number of observations or UKBB participants, *p* is the number of cerebellar subregions, and *q* is the number of whole-brain cortical subregions or cortical networks. Each column of the two data matrices was *z*-scored to zero mean (i.e., centering) and unit variance (i.e., rescaling) across the n participants.

This way, we had four input variable sets for PLSR and thus derived four dedicated PLSR models, each modelling a unique feature composition of cortical granularity and cerebellar modality of cortico-cerebellar covariation (Table 1).

**Table 1.** Four combinations of cerebellar modalities and cortical parcellation granularity yielded four PLSR models.

| Cerebellum \ Cortex | Cortex |  |
| --- | --- | --- |
|  | 100 cortical parcels | 7 cortical networks |
| 28 functionally derived cerebellar parcels | Funct100region PLSR | Funct 7net PLSR |
| 28 structurally derived cerebellar parcels | Struct100region PLSR | Struct 7net PLSR |

The PLSR algorithm then addressed the problem of maximizing the covariance between the low-rank, mutually orthogonal projections from the two variable sets or data matrices. The two sets of linear combinations of the original variables are obtained by PLSR as follows:

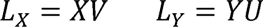

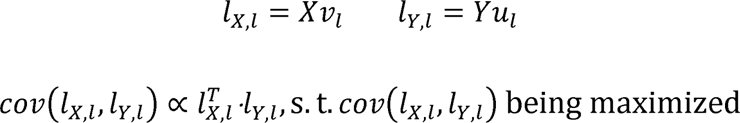

where orthonormal *V* and *U* indicate the respective contributions of *X* and *Y*, *L_X_* and *L_Y_*denote the respective latent “modes” expression of joint variation based on the key patterns derived from *X* and from *Y*, *l_X_*_,*l*_ is the *l*th column of *L_X_*, and *l_Y_*_,*l*_ is the *l*th column of *L_Y_*. Note that for a given latent mode, the overall signs of its weight vectors v_l_ and u_l_ are arbitrary, that is, overall positive and negative values can be flipped for a vector as a whole. We define modes as general principles of population variation in our target neural circuits that can be reliably extracted in brain structure at the population level. The goal of our PLSR approach was to find corresponding pairs of latent vectors *l_X,l_* and *l_Y,l_* that yield maximal covariance in the derived latent embedding space. The data matrices *X* and *Y* were decomposed into *L* components iteratively, where *L* denotes the number of modes to be estimated by the model. Alternatively, PLSR finds the canonical vectors *u* and *v* that maximize the relationship between a linear combination of cerebellar volume ratio expressions (*X*) and a linear combination of brain volume ratio expressions (*Y*). By doing so, PLSR identifies the two concomitant projections *Xv_l_* and *Yu_l_* which optimized co-occurrence between patterns of whole-cerebellum-cortex and whole-cerebrum-cortex variation across participants.

Each identified mode was indicative of the principle cross-association between a constellation of within-subject cerebellar variations, and a constellation of within-subject cortical variations that occur in conjunction with each other at the population level. The set of *L* orthogonal modes are mutually uncorrelated by construction (Wang et al., 2020). They are also naturally ordered from the most important to the least important cortico-cerebellar covariation pattern based on the amount of variance explained between the latent cerebellar and cortical variable sets. Pearson’s correlation between a pair of latent variables is commonly used to quantify the achieved degree of cross-correspondence between cerebellar subregions and cortical subregions or networks for a given mode. The first and strongest mode therefore explained the largest fraction of joint variational effects between combinations of cerebellar subregions and combinations of cortical regions. Each ensuing covariation mode captured a fraction of structural covariation that is not already explained by one of the *L* −1 preceding modes. The variable sets were entered into PLSR after a confound-removal procedure based on previous UK Biobank research (cf. above).

### Robustness of derived modes via empirical permutation testing

To assess the solidity of each identified mode, we applied the identical analysis pipeline (as above) based on 1000 permutation iterations. In each iteration, we constructed empirical noise models by random permutation of the preprocessed cerebellar parcel measures. Once constructed, each permuted variable set was fed into a PLSR model as variable set *X* with the unpermuted, original cortical features (variable set *Y*) and decomposed into *L* candidate cortex-cerebellum covariation modes (as above). Each of our actual modes of PLSR models was then compared with the hypothetical mode whose explained variance was on average the highest across 1000 permutation iterations, which yielded a null distribution of the expected explained variance for that actual mode. We kept only the actual modes that had a p-value < 0.001. This permutation analysis scheme intentionally aimed to break the link between homologous cortex-cerebellum pairs within individuals to generate an empirical null distribution. These noise data simulations allowed testing of the robustness of covariation patterns in the hypothetical population.

### Phenome-wide profiling

To enrich our core analysis (cf. above), we aimed to interrogate the practical implications of the derived cortico-cerebellar covariation modes. We sought to understand their links to UK Biobank traits across a deliberately broad spectrum of predefined phenotype and categories. To that end, we performed an in-depth annotation of the cortico-cerebellar modes by screening 977 phenotypes encompassing lifestyle factors, cognitive tests, and physical health assessments. To carry out this phenome-wide association analysis (PheWAS), we sourced phenotypic features through two specialized tools tailored for aggregating, cleaning and normalizing UK Biobank phenotype data according to predefined rules. We started with a raw collection of ∼15,000 phenotypes that were fed into the FMRIB UK Biobank Normalisation, Parsing And Cleaning Kit (FUNPACK version 2.5.0; https://zenodo.org/record/4762700#.YQrpui2caJ8). Through FUNPACK, we curated and harmonized the data, focusing on a collection of phenotypes linked to 11 categories of interest. The results from FUNPACK, which yielded ∼3,300 phenotypes, were then fed into PHEnome Scan ANalysis Tool (Millard et al., 2018) (PHESANT, https://github.com/MRCIEU/PHESANT) for further refinement, cleaning and data categorization. The resulting final set of 977 phenotypes of interest derived from PHESANT was subsequently analyzed against discovered covariation modes to probe for associations between cortico-cerebellar structural covariation pattern expressions and target phenotypes.

To be specific, we used FUNPACK on the UKBB sample to extract phenotype information covering 11 major categories, including ‘blood assays’, ‘cognitive phenotypes’, ‘early life factors’, ‘lifestyle and environment - alcohol’, ‘lifestyle and environment - exercise and work’, ‘lifestyle and environment - general’, ‘lifestyle and environment - tobacco’, ‘mental health self-report’, ‘physical measures - bone density and sizes’, ‘physical measures - cardiac & blood vessels’, ‘physical measures - general’. These categories of interest were predefined in the FUNPACK utility (-cfg fmrib arguments) and they excluded any brain-imaging-derived information. As there were only four phenotypes associated with diet, we excluded this category from downstream analysis. The FUNPACK setting defining the phenotype categories contained a built-in toolkit tailored to the UKBB which we used to refine the phenotype data, such as removing ‘do not know’ responses and replacing unasked dependent data. For example, a participant who answered that they do not use mobile phones was not asked how long per week they spent using a mobile phone. Here FUNPACK would fill in a value of zero hours per week as a response. As such, FUNPACK’s built-in rule-based pipeline yielded 3,330 high-quality phenotype columns.

The intermediate output from FUNPACK was then fed into PHESANT, which is an established tool for further curating UKBiobank phenotypes. In addition to combining phenotypes across visits, data cleaning and normalization, PHESANT categorized the data as belonging to one of four datatypes: categorical ordered, categorical unordered, binary or numerical. All categorical unordered columns were one-hot encoded into binary columns to represent a single response. For example, the employment status phenotype was originally encoded as a set of categorical indicators representing different contexts (e.g., retired, employed, on disability). These conditions were converted into a binary column (e.g., retired true or false). We then combined the output of categorical one-hot encoding of unordered phenotypes with all measures classified by PHESANT as binary, numerical, or categorical ordered. In so doing, the final set comprised 977 phenotypes.

We deployed both FUNPACK and PHESANT with their default parameter choices. As a result, all columns containing fewer than 500 participants were automatically discarded from further analysis according to PHESANT’s default protocol. Moreover, FUNPACK by default assessed pairwise correlation between phenotypes and kept only one phenotype of a set of highly correlated phenotypes (>0.99 Pearson’s correlation rho). For example, left and right leg fat percentages were highly correlated (Pearson’s r = 0.992). Hence, only right leg fat percentage was included in the final set of phenotypes. The choices of which phenotypes to discard were also automatically streamlined and conducted by FUNPACK.

We then explored and charted the relationships between the subject-wise expression of specific cortico-cerebellar covariation patterns, with their single-subject expression, and the portfolio of 977 phenotypes, ensuring appropriate correction for multiple comparisons. By computing the Pearson correlation between each phenotype and the inter-individual variation in the cortical latent variable and cerebellar latent variable respectively, we discerned both the association strength and the accompanying statistical significance of their relationship. For each identified covariation pattern, we applied two widely established corrections for multiple comparisons to account for the numerous association tests being evaluated in our assay. We first utilized Bonferroni’s correction, adjusting based on the number of phenotypes tested (0.05/977 = 5.11e-5). Additionally, we examined the significance of our correlation strength through the false discovery rate (FDR), another widely accepted method for multiple comparison correction that is more lenient than Bonferroni’s approach. The false discovery rate (Benjamini & Hochberg, 1995) was set as 5% (Miller et al., 2016; Raizada et al., 2008; Sha et al., 2021) and computed for each covariation mode following standard practice (Genovese et al., 2002). To aid in visualization, all phenotype results were color-coded and organized based on category membership as per definitions provided by FUNPACK.

## Results

Previous neuroscience studies have rarely directly investigated the covariation between the entire cerebral cortex and the entire cerebellum in humans (Balsters et al., 2010; Hill et al., 2010; Liu et al., 2022), let alone across a large cohort of subjects. Our central analysis yielded different numbers of statistically significant derived latent modes: 2 modes for ‘Funct 7network’, 6 modes for ‘Funct 100region’, 4 modes for ‘Struct 7network’ and 7 modes for ‘Struct 100region’ (Table 1). Here we first attend to the selected cortical and cerebellar structural variation weights (U and V, representing the respective contributions of cortical and cerebellar subregions to the derived latent modes, cf. Methods). We then present the significant phenotype profiles associated with cortical latent variable and cerebellar latent variable.

### Dissociation between higher and lower neural system in cortex-cerebellum covariation

Across examined models (Table 1), the most explanatory and robust sources of population variation revealed a highly consistent higher-level cognitive system and the low-level visual-sensorimotor system interplay in the cortex-cerebellum covariation, no matter which atlas parcellation definition was used. We start with the model ‘Funct 7net’.

Mode 1 exhibited the highest explained variance of 0.41, and juxtaposed a sensorimotor-visual network against higher-order cognitive networks, as manifested by the opposing directions of structural variation trends. As can be seen for the cerebellum (Fig. 1), we observed the positive weights in regions D2 distributed around the border between lateral lobule VI and crus I, between lateral crus II and lobule VIIb, and scattered in lobule IX; generally known to be associated with executive functions (King et al., 2019; Nettekoven et al., 2023). Central and lateral crus I, crus II and paravermal IX, related to the fused social-language regions S2 and S3, also exhibited strong variation weights. The motor associative atlas region M2 related to the mouth at the intersections of bilateral paravermal lobules VIIb and VIIIa showed positive variation weights, too. For the cerebral cortex, we observed structural variation in the executive-control network (ECN), default mode network (DMN), limbic network, and salience / ventral attention network (VSN) (Fig.1A and S1 Fig A). Together, higher-order cognitive function-related regions in both the cerebellum and cortex showed the same direction of structural variation.

**Figure 1.**
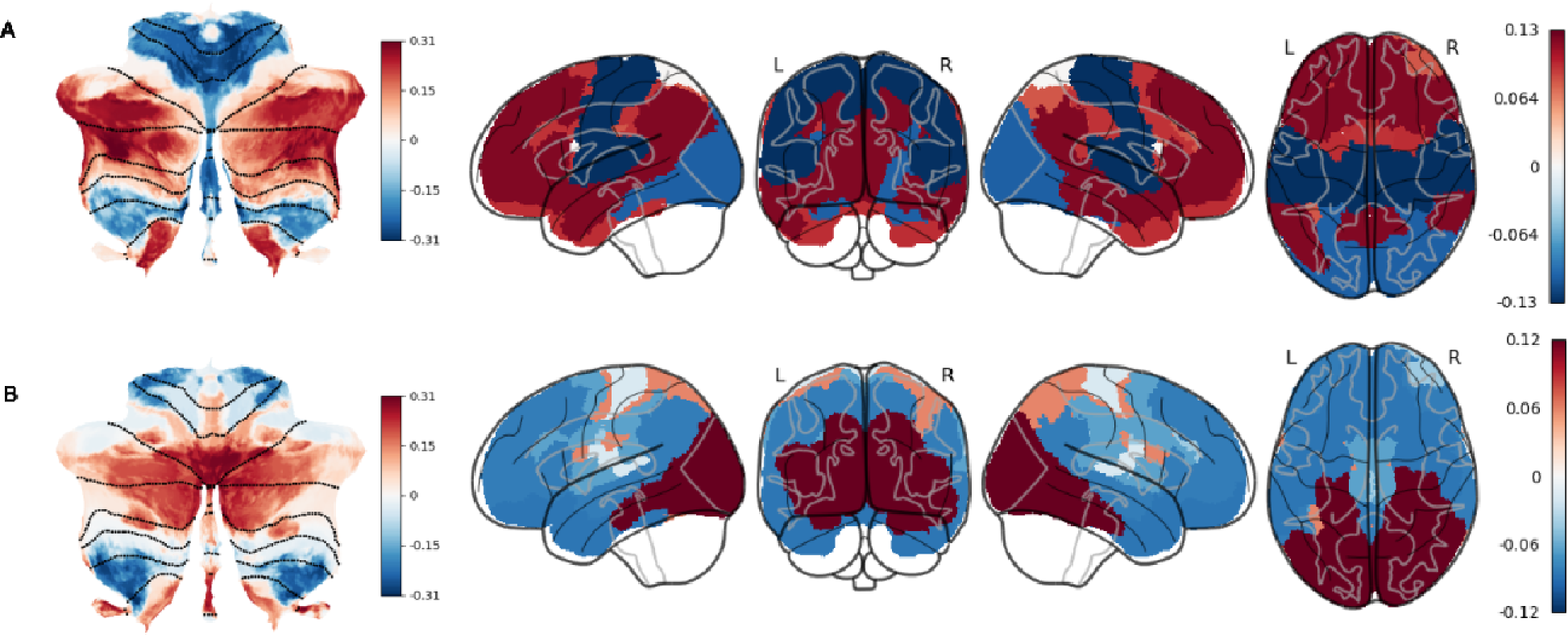
Funct7net Modes 1 and 2 explained contrasting variation in the higher cognitive-lower visuosensorimotor system interplay versus exogenous visuoattentional-endogenous cognitiv system. Mode 1 and Mode 2 in cerebellar functional parcellation with cortical 7 network averaged (‘Funct 7net’) displayed a clear contrast between higher cognitive system and lower visual-somatosensor system. (A) Mode 1 had an explained variance of 0.41. One end of structural variation weights was foun in regions associated with executive functions and social-language processing in the cerebellum an general higher-order cognitive networks in the cerebral cortex. The opposite end of structural variatio weights was observed in other cerebellar regions and the sensorimotor and visual networks in th cerebral cortex. Weak weights were assigned to the Dorsal Attention Network (DAN). The cortical variation patterns align with the DMN-visuomotor distinction, while the cerebellar variation pattern correspond to the classical double motor representation and the triple nonmotor representation. (B) Mod 2 had an explained variance of 0.28. One end of structural variation weights was observed in bilateral ey atlas regions in the cerebellum, as well as in executive-function and social-language regions. Th cerebral cortex exhibited strong variation weights in the visual network and lesser variation weights in th DAN. The opposite end of structural variation weights was found in specific cerebellar regions associate with introspection, action observation, and lower limb functions, which showed structural covariation wit other higher-order cognitive networks in the cerebral cortex. Notably, an antagonist relationship observe between the DAN and other higher-order cognitive networks was potentially anchored in the cerebellum. Red, blue and white indicate positive, negative and zero weights that maximize the covariance between cortical and cerebellar latent variables.

Conversely, in mode 1, sensorimotor and action-related regions of both the cerebellum and the cortex exhibited a structural covariation pattern that was directionally opposite to that observed in the higher-order cognitive regions. For the cerebellum, we observed the strongest negative weights in the upper limbs atlas regions M3 which were defined by a significant proportion of lateral lobules I-V and VIIIb and complex action atlas regions A1 which sit across lobules I-V vermis and extended into lobule VI and further extended from crus II to lobule VIIIb vermis. Additionally, we observed weaker weights in action observation atlas regions A2 sitting on the border between lateral lobules V and VI, between lobules VIIIa and VIIIb and lower limbs atlas regions M4 occupying lateral lobules I-V and VIIIb regions in the cerebellum. Finally, we observed the weakest variation weights in the oculomotor vermis M1 (Nettekoven et al., 2023), which was mostly located in VI vermis, and some portions of lobule IX/X vermis. For the cerebral cortex side of this population mode, we obtained strong contribution in the sensorimotor network and visual network (Fig.1A and S1 Fig A). The cortical weights in mode 1 were broadly consistent with the DMN-visuomotor divergence of neural systems described by Margulies et al. (2016) and, before, by Mesulam (1998) such that the DMN and the unimodal visual, sensorimotor networks’ weights were at the two extreme ends while other higher-order cognitive networks were situated in between. The cerebellar weights in mode 1 were also highly consistent with the most dominant functional gradient in the cerebellum (Guell, Schmahmann, et al., 2018) and with the classical double motor representation (lobules I-VI and VIII) and the recently proposed triple nonmotor representation (lobules VI/crus I, crus II/VIIb, IX/X) (Buckner et al., 2011; Guell, Gabrieli, & Schmahmann, 2018).

Mode 2 showed the second-highest explained variance of 0.28 in our analyses based on the functionally derived cerebellum atlas. Mode 2 separated a visual-attention network from other functional networks which were assigned opposite structural covariation weights. In the cerebellum, we observed one extreme pole of structural variation weights in M1 (cf. above). Executive-function region D1 also exhibited strong implications in mode 2. This region consisted of 2 patches on the cerebellar cortex at the border between central lobule VI and crus I, between central crus II and VIIb. It also stretched along paravermal lobules VI, crus I and crus II. We further observed relevant positive weights in two social-language atlas regions: S1 defined by extensive coverage of paravermal crus I, crus II and areas into lobule VI, VIIb and S2_3 (cf. above). For the cerebral cortex, we observed the most prominent structural variation weights in the visual network and some implications of the dorsal attention network (DAN) (Fig.1B and S1 Fig B).

The opposite direction of structural variation in mode 2 was found in lateral lobule VIIIb and IX, in the fused regions A3 and S5, related to introspection and action observation (Nettekoven et al., 2023). We further observed significant structural variation weights in lower limbs regions M4 and action observation region A2 (cf. above). These cerebellar regions showed systematic structural covariation with default mode network (DMN), limbic network, VSN and executive control network (ECN) in the cerebral cortex (Fig.1B and S1 Fig B). Notably, our findings witness an antagonist relationship between DAN and other higher-order neural systems here modeled as anchored in the cerebellum, which was less recognized than their established anticorrelation relationship in the cortex (Andrews-Hanna et al., 2014; Brissenden et al., 2016; Fox et al., 2005; Spreng, 2012).

### 100-region cortical parcellation mirrors the dissociation in 7-network cortical atlas

Our second model used cerebellar functional parcellation with cortical 100 region parcellation (‘Funct 100region’, Table 1) which is based on the same cortical atlas but differed from the ‘Funct 7net’ model (cf. above) by higher resolution. The left and right homologous regions in 100 region parcellation are also separated, while in the 7 network parcellation there is no left-right hemispheric distinction. Our ‘Funct 100region’ model broadly reiterated the trend of motor vs cognitive systems detailed in mode 1 (explained variance 0.49, Fig 2A and S2 Fig A) and visuoattention network’s anticorrelation with other network systems in mode 2 (explained variance 0.32, Fig 2B and S2 Fig B) with the more granular cortical subregional specification. The cerebellar weights in each pair of mode 1 and mode 2 of ‘Funct 100region’ and ‘Funct 7net’ were quite similar, with mode 1 absolute Pearson’s |r| = 0.99, p-value < 10^-29 and mode 2 absolute Pearson’s |r| = 0.98, p-value < 10^-19, after correlating across respective cerebellar region weights of the 2 cerebellum-brain covariation model instances.

**Figure 2.**
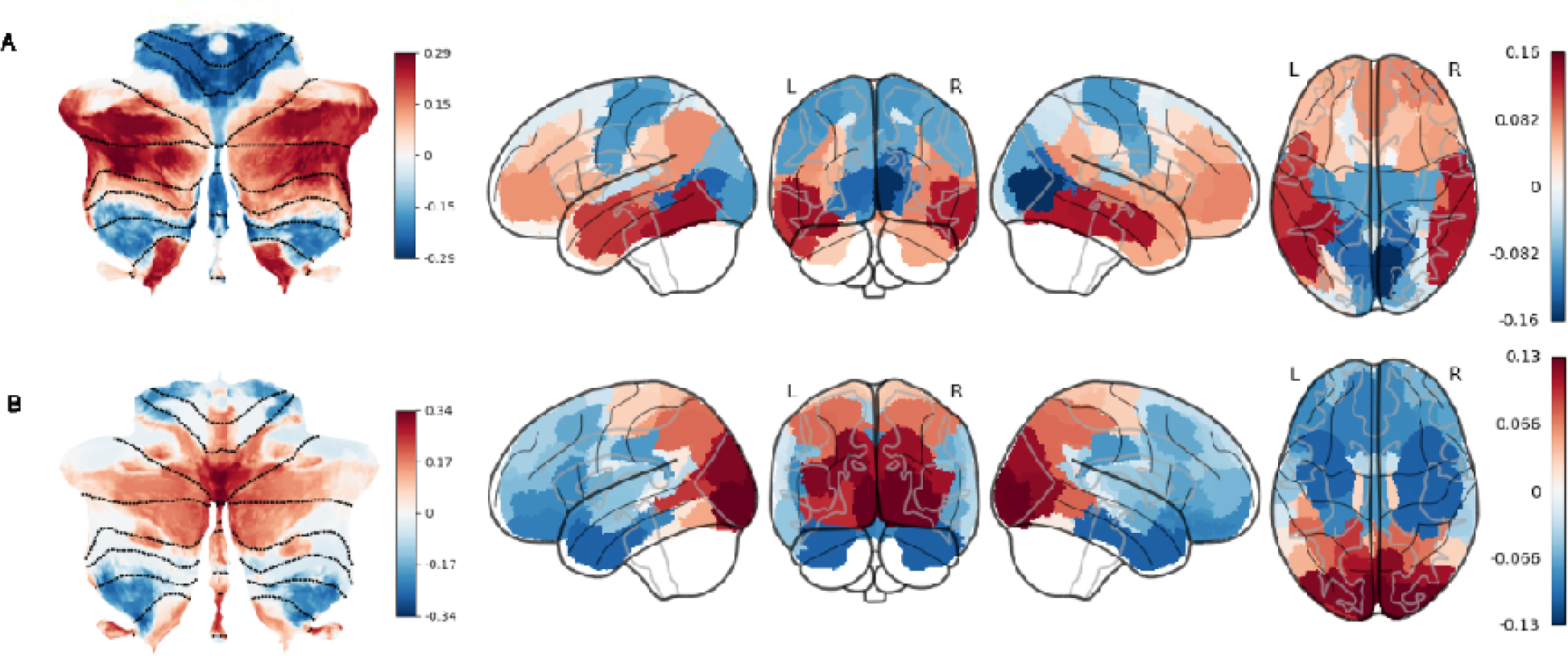
Funct100region Modes 1 explains opposite variation in primary visual / sensorimotor regions versus higher associative regions, while mode 2 explains visual-attentional regions to anticorrelate with higher associative regions. Mode 1 and mode 2 in cerebellar functional parcellation with cortical 100 region parcellation (‘Funct 100region’) broadly reiterate the patterns observed in the first two modes of ‘Funct 7net’, with almost perfectly correlated cerebellar weights respectively (mode 1: r = 0.99, p-value < 10^-29; mode 2: r = 0.98, p-value < 10^-19). As cortical weights were more different, here we only describe the differences in cortical weights. For the description of cerebellar weights, see Figure 1. (A) mode 1 had an explained variance of 0.49. One end of structural variation weights wa concentrated in the bilateral temporo-occipital cortex. The opposite end of structural variation weights was distributed in bilateral parieto-occipital lobes and the primary somatosensory cortex. (B) mode 2 had a explained variance of 0.32. One end of structural variation weights was observed in bilateral parieto-occipital lobes. The opposite end of structural variation weights was found in bilateral frontal cortex an anterior temporal lobe.

While the pattern of cerebellar weights was extremely similar, the ‘Funct 100region’ solutions provided a more fine-grained analysis of the cortical patterns than the ‘Funct 7net’ solution showing that the structural covariation concentrates on certain subsets of regions within each network rather than on the entire network. For mode 1, one end of the structural variatio weights was concentrated in bilateral temporo-occipital cortex including left visual word form area, left middle-inferior temporal gyrus, left temporal pole, right middle-inferior temporal gyrus and right fusiform gyrus. On the other end of structural variation were bilateral parieto-occipital lobes including primary visual cortex, extrastriate cortex, retrosplenial cortex, lingual gyrus, primary sensorimotor cortex, premotor cortex and surprisingly primary auditory cortex (Fig 2 and S2 Fig A). In mode 2, we observed one extreme end of structural variation weights i bilateral parieto-occipital lobes including primary visual cortex, extrastriate cortex, fusifor gyrus, intraparietal sulcus (IPS) and cuneus, retrosplenial cortex and precuneus, and the other end of structural variation weights in bilateral frontal cortex and anterior temporal lobe, including the temporal pole, parahippocampal gyrus, orbitofrontal cortex, ventrolateral prefrontal cortex and anterior insula cortex, cingulate cortex, dorsolateral prefrontal cortex (Fig 2B and S Fig B).

### An ipsilateral axis of cortical-cerebellar variation

Despite the known mainly contralateral fiber connection between the cerebellum and the cerebral cortex, we persistently observed an ipsilateral pattern in the 100-region cortical parcellations such that the dominating regions in the cerebellum and cerebral cortex variation share the same end of weights for a given side of the brain.

Mode 3 (‘Funct 100region’) showed an explained variance of 0.30 (Fig.3 and S2 Fig C). Notabl, the entire left and right cerebellum showed opposite variation trends. In the left cerebellum, w observed the most significant structural variation concentrated in regions centering on the posterior lobule VI, lateral anterior crus I, lateral VIIb, anterior VIIIa, lobule IX / X and vermal I-VI that are associated with executive function (working memory) as per atlas, areas of paravermal crus I, crus II and areas extending into lobule VI, VIIb that are related to social-language and the intersections of bilateral paravermal lobules VIIb and VIIIa related to mouth i the left cerebellum. In this mode’s counterpart in the cortex, left cortical regions varied in th same direction as the left cerebellum, especially the primary auditory cortex, Broca’s area, lower primary sensorimotor regions and inferior frontal gyrus. On the contrary, the right cerebellum varied in the same direction as right cortical regions, especially the right cerebellar regions associated with executive function, social-language, motor and complex action; right cortical regions, such as inferior/middle temporal gyrus, fusiform gyrus, TPJ and primary auditory corte . Interestingly, the cortical regions conforming to the usual contralateral patterns were th precuneus and posterior cingulate cortex, albeit with less significant weights. Together, mode (‘Funct 100region’) displayed a robust ipsilateral pattern such that either side of cerebellum varied in the same direction as the dominant cortical regions on the same side of the brain.

**Figure 3.**
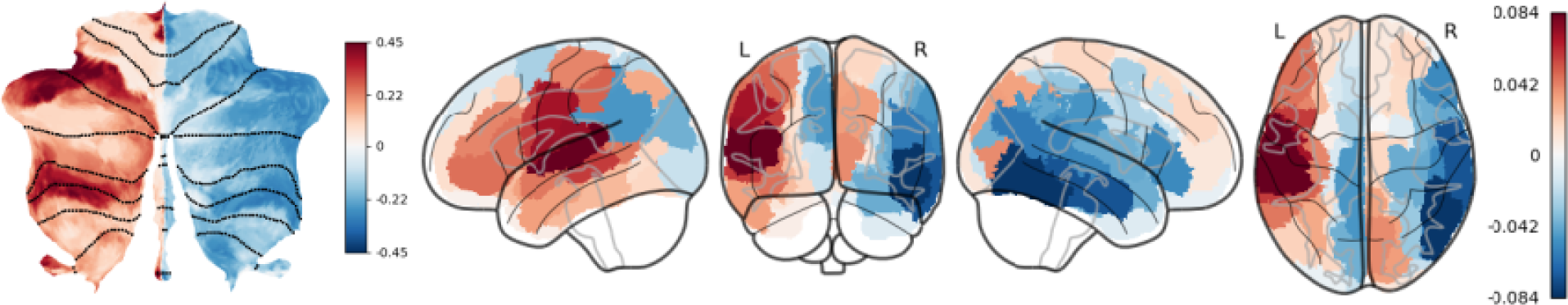
Driving ipsilateral principles in cerebellum-cortex correspondence as a key feature of population brain variation. An ipsilateral pattern for the dominating weights’ direction in the cerebellum and cerebral cortex appeared in ‘Funct 100region’ mode 3. (A) ‘Funct 100region’ mode 3 had an explained variance of 0.30. The entire left cerebellum and right cerebellum showed completely opposite directions of structural variation weights. One end of structural variation weights concentrated in specific regions associated with executive function, social-language, and motor functions in the left cerebellum, and in the left cerebral hemisphere including the primary auditory cortex, Broca’s area, and inferior frontal gyrus. Conversely, the other end of structural variation weights observed in right cerebellar region roughly symmetric to the left side, associated with executive function, social-language, and motor functions, and in the right cerebral hemisphere including inferior/middle temporal gyrus, fusiform gyrus, TPJ, and primary auditory cortex. The precuneus and posterior cingulate cortex exhibited a contralateral pattern.

The ipsilateral cerebellar-cortical pattern was also observed in a mode based on the structural atlas: mode 2 (‘Struct 100region’; S4 Fig A and S8 Fig A) which had an explained variance of 0.29, on par with mode 3 (‘Funct 100region’). One end of the most prominent structural variation weights was seen in right crus I, crus II and left lobule VIIIa/b, and in the right superior temporal sulcus, auditory cortex, somatosensory cortex, TPJ, dlPFC; the other end of the most significant structural variation weights were seen in left crus I, crus II and right lobule VIIIa, and in the left temporal lobe, fusiform gyrus, supramarginal gyrus, angular gyrus and some visual/auditory/motor-associative areas. Again, the only cortical regions conforming to the usual contralateral patterns were the precuneus and posterior cingulate cortex.

Comparing the cerebellar weights in modes derived from the functional vs. structural cerebellum atlas, the cerebellar weights in the former were more homogeneous for either side such that all the cerebellar regions on the left show the opposite direction of structural variation to all the cerebellar regions on the right, while the latter showed both directions of structural variation on each side. Comparing mode 3 (‘Funct100region’) and mode 2 (‘Struct100region’), we found that the cortical weights were highly similar (Pearson’s r = −0.82, p-value < 10^-25). Thus, this suggests that the modes from the functionally and structurally derived cerebellar atlas point to at least some overlapping associative brain findings. Overall, the strongest weights still followed the ipsilateral pattern.

### Anatomical cerebellar regions capture less structural variation

We focused our results on the functionally derived cerebellum atlas because the explained variance of mode 1, mode 2, mode 3 and mode 4 from the structurally derived cerebellum atlas were 0.20, 0.19, 0.13 and 0.12 respectively, achieving only about half the explained variance of those from the functionally derived cerebellum atlas. This observation suggested that the functionally derived cerebellum atlas explained the data better. We also found these modes were harder to interpret, especially regarding the cerebellar latent variables. But these cerebellar patterns arguably reconciled with several classical cerebellar functional organizations, such as Crus I/II’s crucial role in DMN and attention, lobule I-IV / VIII’s role in motor function and vermis or flocculonodular lobe’s role in limbic function (Chen et al., 2023; Schmahmann, 1991).

Struct 7net’s mode 1 (S3 Fig A and S7 Fig A) and mode 4 (S3 Fig D and S7 Fig D) generally reproduced a similar pattern to those of ‘Funct 7net’ mode 1’s (Fig. 1A and S1 Fig A) cortical weights after correlating across respective cortical regions, with mode 1 absolute Pearson’s |r| = 0.94, p-value < 10^-47 and mode 4 absolute Pearson’s |r| = 0.92, p-value < 10^-42. In mode 2 (S3 Fig B and S7 Fig B) of ‘Struct 7net’, we captured a strong reciprocal relationship between DAN and DMN / limbic network that was highlighted in mode 2 of ‘Funct 7net’ (Fig.1B and S1 Fig B), which added more evidence that an antagonist relationship between DAN and DMN could be anchored in the cerebellum. In mode 3 of ‘Struct 7net’ (S3 Fig C and S7 Fig C), we observed a similar anti-correlation between ECN-DAN and DMN-limbic system. For mode 1, the cerebellar regions varying concurrently with the higher order cognitive networks except DAN were bilateral crus I/II and lobule X, whereas the cerebellar regions consistent with sensorimotor network were lobules VIIIa/b, lobules IX and anterior lobe to a lesser extent. This pattern roughly forms a double-motor (lobules I-VI, VIII and additionally IX), triple-nonmotor representation (lobules crus I, crus II and X) in the cerebellum (Guell, Gabrieli, & Schmahmann, 2018). For mode 2, the cerebellar regions tied with DMN and limbic networks were anterior lobe, vermis across the posterior lobe, bilateral lobules VIIIb and lobule X, whereas the cerebellar regions coherent with DAN were bilateral lobules VII. For mode 3, the cerebellar regions covarying with the ECN and DAN were lobule X vermis, anterior lobe, bilateral lobules VIIIb and IX and Crus I, while the cerebellar regions covarying with DMN and limbic system were bilateral lobules VIIb, Crus II and lobules VIIIa. For mode 4, the cerebellar regions coherent with the higher-order cognitive networks except limbic network were the posterior lobe up to crus II, excluding most of the cerebellar vermis or lobule X, while the cerebellar regions consistent with visual-motor networks were the anterior lobe plus crus I to a lesser extent, which reminded us of the classical anterior motor - posterior cognitive cerebellum segmentation (Schmahmann, 2004; Schmahmann et al., 2019; Schmahmann et al., 2009; Schmahmann & Sherman, 1998; Stoodley & Schmahmann, 2009b; Stoodley et al., 2012).

Finally, mode 3 (S4 Fig B and S8 Fig B) in our structurally derived cerebellum atlas with cortical 100 region parcellation (‘Struct 100region’) corresponded to ‘Funct 100region’ mode 1 (Fig 2A and S2 Fig A), and their cortical weights were highly correlated with absolute Pearson’s |r|=0.85, p-values < 10^-27. It also corresponded to ‘Struct 7net’ mode 1 (S3 Fig A and S7 Fig A), with highly correlated cerebellar weights reaching absolute Pearson’s |r|=0.93, p-values < 10^-11. Mode 5 (S4 Fig C and S8 Fig C) corresponded to ‘Funct 100region’ mode 2 (Fig 2B and S2 Fig B) and ‘Struct 7net’ mode 2 (S3 Fig B and S7 Fig B), with an explained variance of 0.21. The cerebellar weights between mode 5 and ‘Struct 7net’ mode 2 were highly correlated with Pearson’s r=0.84, p-values < 10^-7. However, the cortical weights were very different from ‘Funct 100region’ mode 2. We observed the frontal eye field and IPS which are key regions of the DAN dominated one end of variation, while RSC, insular cortex, auditory cortex and inferior prefrontal cortex, which play important roles for episodic memory, limbic function and linguistic function dominated the other end. This pattern overall corresponded with the DAN vs. DMN-limbic contrast seen in Struct7net mode 2’s cortical weights. The explained variances of 0.27 and 0.21 were still much less than the explained variance of ‘Funct100region’ mode 1 and 2, which reaffirmed our focus on the analysis of functionally derived cerebellum atlas.

### Funct7net mode 1 shows phenotypic signature with distinct arrays of cognitive, lifestyle, physical conditions, and blood traits

To understand relevant real-world implications of the derived cerebellar-cortical variation modes, we carried out phenome-wide association assays (PheWAS): We tested links of 977 phenotype measurements across all the subjects with cerebrocerebellar modes 1 and 2 from Funct 7net (Fig.4, 5) and Funct 100region PLSR results (S5,6). This revealed the unique phenotypic profiles of each mode and provided a wealth of potentially undiscovered brain-phenotype connections. We paid special attention to the phenotypes whose association with either latent variable was above the Bonferroni correction thresholds.

**Figure 4.**
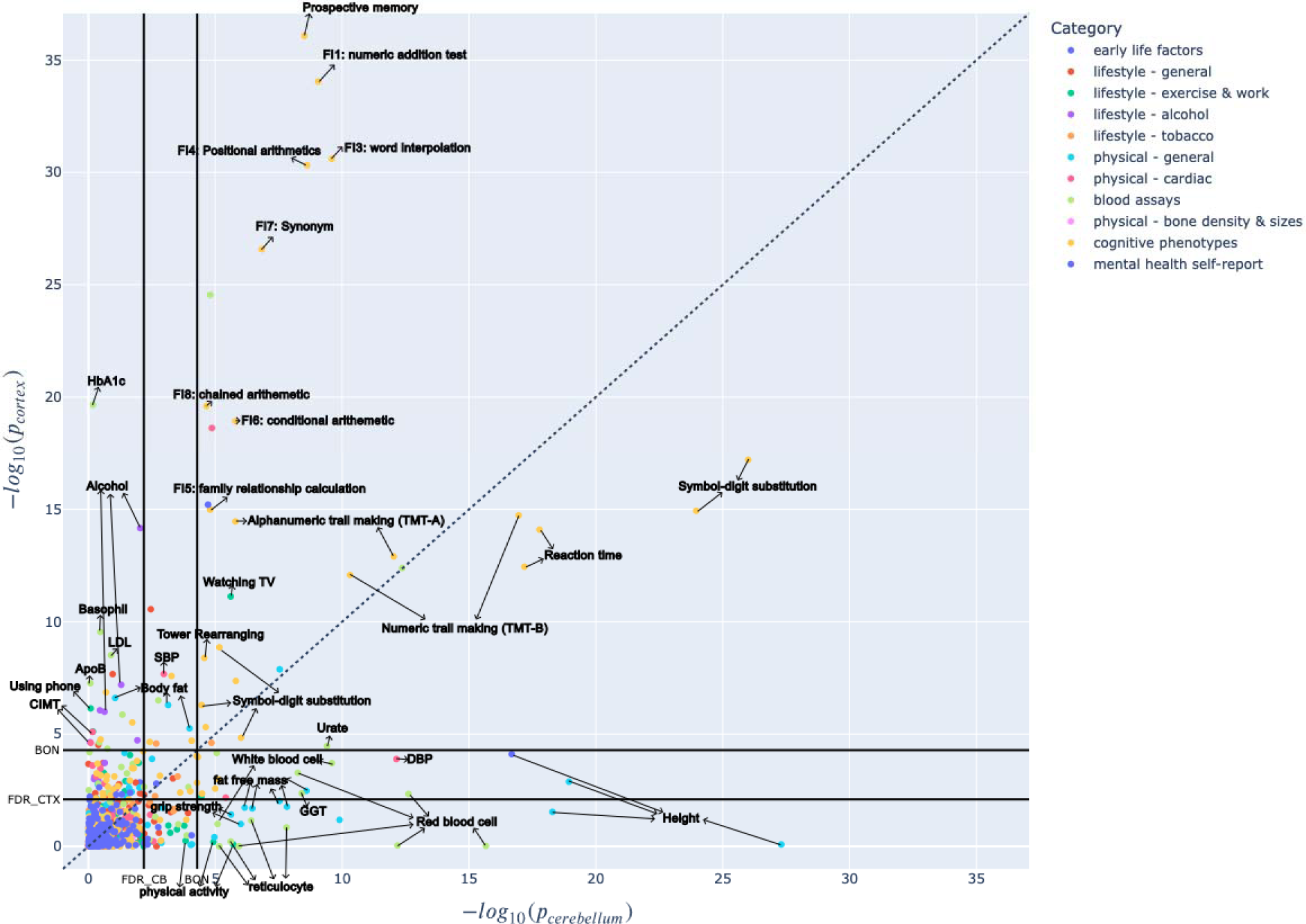
Funct 7net Mode 1 shows phenotypic signature with distinct arrays of cognitive, lifestyl, physical conditions, and blood traits for the higher-lower system correspondence across th cortex and cerebellum. Mode 1 (Funct7net) cerebellar-cortical phenotype profiles. The x-axis showe the −log_10(p-value) of the cerebellum while the y-axis showed the −log_10(p-value) of the cortex. Th Bonferroni correction threshold (BON) and False Discovery Rate (FDR) are plotted for the cortex and th cerebellum on the respective axis. Phenotypes near the upper-left part of the figure are more strongl associated with the cortex while those near the lower-right corner are more strongly associated with th cerebellum. The phenotypes near the diagonal are relevant for both the cortex and the cerebellum. Th phenotypes at the lower-left corner bordered by the two BON threshold lines are considered insignificant.

In mode 1 (Funct7net; Fig. 4), where the higher cognitive - lower sensorimotor system interplay explained the most cerebello-cortical covariance, overall, there appeared to be a closer link between complex cognitive ability, obesity, the likelihood of cardiovascular diseases, a sedentary lifestyle and the cortical latent variable. Additionally, psychomotor processing speed, physical activity and angiogenesis were better associated with cerebellar latent variable.

The association between many cognitive phenotypes and the latent cortical and cerebellar variables surpassed other phenotype categories by a margin. Generally, most of the significant cognitive phenotypes are common to both the cortex and the cerebellum. Yet, they showed more dominant associations with the cortex component than with the cerebellum component of our mode 1, including mostly fluid intelligence tests, alphanumeric trail-making tests (TMT-B), initial answer for prospective memory and matrix completion tasks that require abstract reasoning, complex executive functions, focused visual attention and motor control (Beaty et al., 2015; Beaty et al., 2014; Benedek et al., 2014). The phenotypes that showed equivalent or even stronger associations with the cerebellum are digit symbol substitution test (DSST) (Wechsler, 1944), reaction time test and numeric trail-making test (TMT-A), a widespread hand-eye coordination test, which measures the psychomotor speed and overall efficiency of operations and are known to be closely related to the cerebellum, as evidenced in previous studies (Bolcekova et al., 2017; Botez et al., 1989; Leiner et al., 1989; Stoodley & Schmahmann, 2009a). Nonetheless, both the cerebellar and cortical mode 1 (Funct7net) showed robust associations with a number of cognitive phenotype measures for psychomotor speed, general cognitive ability, cognitive control, response inhibition, task switching, spatial attention, non-verbal reasoning and fluid intelli-gence.

Additionally, we identified a wide range of physiological, physical and behavioral phenotypes distinct for either the cortex or the cerebellum latent variables. Walking and other vigorous physical activities yielded more significant p-values for the cerebellum, but time spent watchin TV and length of phone usage, which are both sedentary behaviors that add to the risks of weight gain and obesity, were more related to the cortex. Interestingly, the phenotypes uniquely linked to the cerebellum, as opposed to the cortex, are largely general physical properties such as height, fat-free mass (FFM) and hand grip strength. Instead, measurements of body fat (or fat mass) are unique to the cortex. Diastolic blood pressure (DBP) and ventricular rate (heart rate) were uniquely associated with the cerebellum but systolic blood pressure and carotid intima-medi thickness which is a measure used to diagnose the extent of carotid atherosclerotic vascular disease were related to the cortex. Finally, alcohol intake was solely associated with the cortex.

In fact, the blood traits also displayed a coherent link with variation in the cortex and for the cerebellum separately. For the cerebellum, several red blood cell-related phenotypes are highly significant (e.g., haemoglobin concentration, mean corpuscular volume/haemoglobin/concentration, hematocrit percentage). Other associated phenotypes included urate (or uric acid) related to kidney function, monocyte and leukocyte (both are white blood cell) count, gamma-glutamyl transferase (GGT, related to the liver), reticulocyte (immature red blood cells) count and percentage. On the contrary, the phenotypes observed only for the cortex included glycated haemoglobin (HbA1c, diabetes), basophil (a white blood cell mainly participating in allergic and inflammatory reactions) count, LDL (low-density lipoprotein-cholesterol or ‘bad’ cholesterol) and Apolipoprotein B (ApoB, cardiovascular disease). This observation also seemed to go hand in hand with more physical activity associated with the cerebellum, therefore increased angiogenesis and blood cells (Isaacs et al., 1992), but more sedentary behaviours in the cortex, which is associated with increased risk of diabetes (Bellettiere et al., 2018; Hamilton et al., 2014), obesity and cardiovascular diseases. Interestingly, insulin growth factor 1 (IGF-1) showed equal association with the cortex and the cerebellum.

### Funct7net mode 2’s phenotype array is driven by time spent watching TV

In mode 2 (Funct7net, Fig 5), which separates the visual attention system from other higher-order neural systems, the driving phenotype for the cortex is time spent watching TV which surpassed other phenotypes by magnitudes. It is also among the top phenotypes associated with the cerebellum. Sleep-related phenotypes such as sleep duration and nap during the day were more related to the cortex than the cerebellum while snoring was more related to the cerebellum. Alcohol was mostly related to the cortex but also has a significant link with variation in the cerebel-lum pattern.

**Figure 5:**
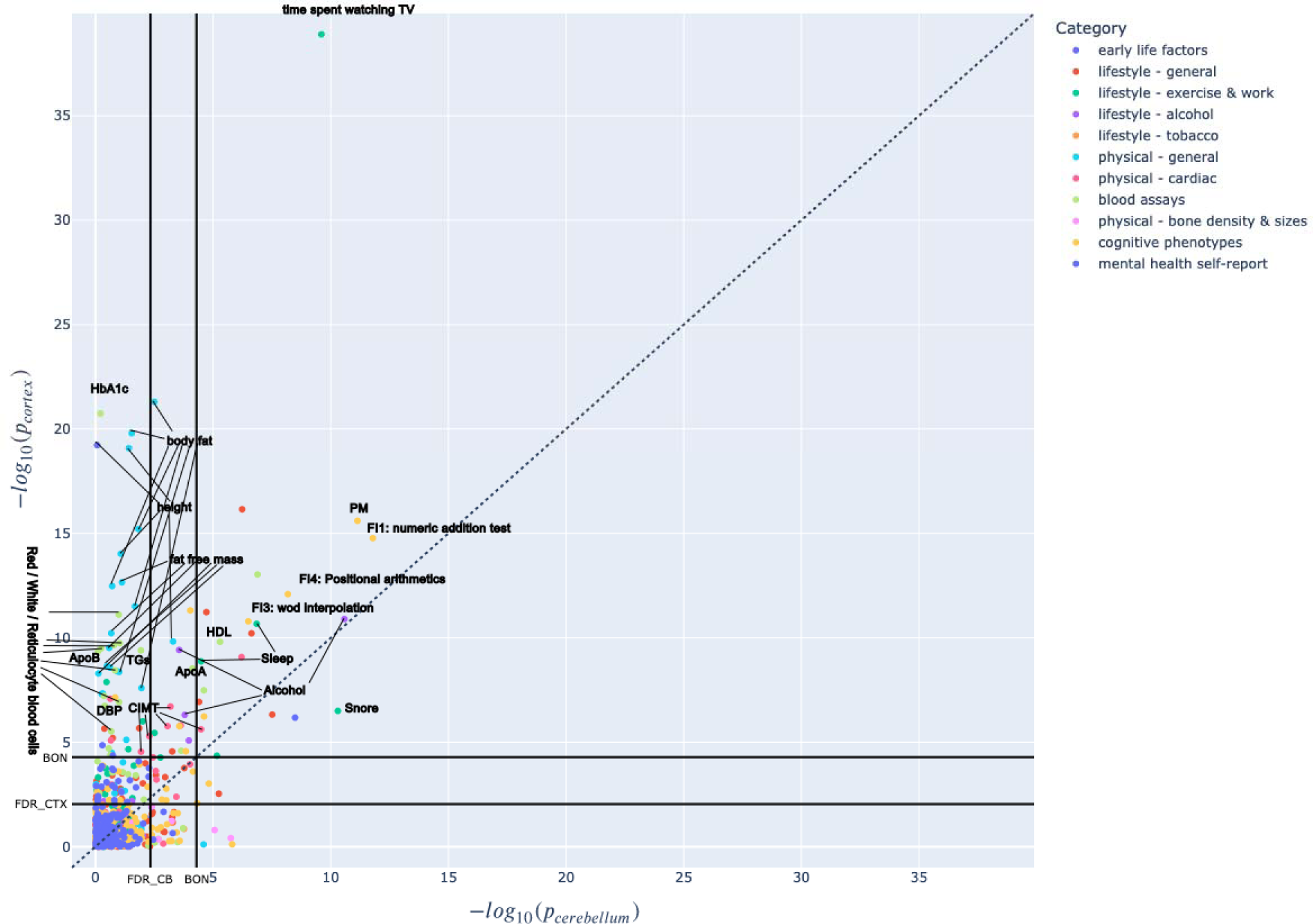
Funct7net mode 2’s phenotype array is driven by time spent watching TV. Most phenotypes show more unique associations with the cortex than the cerebellum, but cognitive phenotypes for fluid intelligence and executive function are significant for the cortex and the cerebellum. The x-axis showed the −log_10(p-value) of the cerebellum while the y-axis showed the −log_10(p-value) of the cortex. The Bonferroni correction threshold (BON) and False Discovery Rate are plotted for the cortex and the cerebellum on the respective axis. Phenotypes near the upper-left part of the figure are more strongly associated with the cortex while those near the lower-right corner are more strongly associated with the cerebellum. The phenotypes near the diagonal are relevant for both the cortex and the cerebellum. The phenotypes at the lower-left corner bordered by the two BON threshold lines are considered insignificant.

The cognitive phenotypes that were significant were again fluid intelligence with a focus on linguistic and arithmetic ability (FI1: numeric addition test, FI4: positional arithmetic, FI3: word interpolation, FI7: synonym, FI6: conditional arithmetic) and initial answer for prospective memory. Their associations with the cortex and the cerebellum were more equivalent than in mode 1.

Further, the BMI, height, blood assay and cardiac phenotypes were more uniquely related to th cortex, possibly because the mode 2’s cerebellar latent variables only emphasized some regions (Funct7net, Fig.1B) instead of the entire cerebellum. It is also possible that even if some phenotypes showed more association with the cortex or the cerebellum across modes, these phenotypes were related to both the cortex and cerebellum. For example, the haemoglobin concentration and haematocrit percentage could be linked to both the areas in the cortex and the cerebellum. Thus, we do not discuss them again.

### Most brain-trait links are congruent between Funct7net and Funct100region mode 1 and 2

Given the similarity of modes between Funct7net and Funct100region, especially for the cerebellar variation, we expected to see a similar phenotype signature in Funct100region mode 1, 2 (S5 and 6) compared to Funct7net mode 1, 2 (Fig. 4 and 5). Indeed, most phenotypes were congruent.

However, for the cortical latent variable of Funct100region mode 1, we observed several differences. First, the cortical association with IGF-1 was the strongest. HbA1c, basophil count and LDL were uniquely associated with the cortex, but the associations were much smaller. IGF-1 has an important role in brain development, neuroplasticity and neuroprotection after brain injury. Expiratory measurement (peak expiratory flow [PEF] and forced expiratory volume in 1 second [FEV1]) were more related to the cortex than the cerebellum, possibly because of oxygenation in brain structure and function. Systolic blood pressure and carotid intima-media thickness measures were insignificant.

Finally, time spent watching TV was associated more with the cerebellum than the cortex, along with physical activity (walking, climbing stairs), which was not observed in Funct7net mode 1. Phone usage was also not significant. Because the cortical latent variable in Funct100region mode 1 emphasized mostly the associative regions between the sensorimotor system and higher-cognitive system, the cognitive phenotypes were uninterrupted.

### Funct100region mode 3 phenotype profiles’ link to ipsilateral brain pattern were hidden

In Funct100region mode 3 where we see the ipsilateral cerebellar-cortico correspondence, most significant phenotypes showed stronger associations with the cortex than with the cerebellum. These significant phenotypes include TMT-A/B, number of siblings, ventricular / pulse rate, reticulocytes counts / percentages, tobacco smoking, IGF-1, handedness, FI3:Word interpolation, corpuscular haemoglobin as well as matrix completion – an aspect of IQ testing (S9 Fig).

## Discussion

### Overview

The cerebellar and cerebral cortex closely interact in supporting everyday activities, from body movement to solving complex problem solving (Ito, 2008; Kelly & L., 2003; Schmahmann et al., 2019; Strick et al., 2009). Yet, it remains mysterious how different parts of these two structures work hand in hand with each other (Chen et al., 2023; Zhu et al., 2023). Here, we designed a mission-tailored analytical framework to 38,527 UKBB participants’ structural brain scans to analyze multiple cerebellum-cortex population covariation patterns and understand how these patterns relate to 977 phenotypic measurements. The most explanatory population mode exhibited the anti-correlation between higher-order cognitive networks (except dorsal attention network) and low-level visual-sensorimotor network within the cerebellum, which mirrors the cortex. The second mode revealed a contrast between the visual attention system and other higher order cognitive system, especially the DMN and limbic system. Another mode revealed an interesting but puzzling ipsilateral structural covariation pattern, despite the mainly contralateral ties between the cerebral and cerebellar cortex. Finally, the phenotypic profiles of the most explanatory patterns unveiled an array of real-world implications with mind, body and exposure of everyday life.

One important finding of this paper was that the functional parcellation explains the structural covariation across individuals better than structural parcellation, especially in the most explanatory modes. This is somewhat surprising, as regions in the functional parcellation often are spatially discontinuous, consisting of multiple parts. If anatomical variation was simply spatially correlated, the more contiguous anatomical parcellation should have performed better. The results imply that not only functional boundaries (King et al., 2019), but also anatomical variation, are better described using functionally defined regions, and that there is value in not respecting lobular microanatomical boundaries.

The uniform cytoarchitecture across the whole cerebellar cortex is well-established (Buckner, 2013; Cajal et al., 1995; Itō, 1984; Voogd & Glickstein, 1998). Despite what appears like an anatomical fact, it has been suspected before that circumscribed territories in the cerebellum have privileged relationships to specific parts of the brain (Schmahmann, 1991, 1996, 1998, 2000; Schmahmann et al., 2019; Schmahmann & Sherman, 1998). Even more so, the cerebellum may harbor a homotopic map of the cerebral cortex (Buckner et al., 2011). Our findings confirm and extent this contention. Additionally, we recontextualize the increasing recognition of a cortical high-low gradient of neural systems given that specific parts of the cerebellum reliably mapped to both low-level sensorimotor and high-level cognitive cortical systems.

### Visual attentional representation in the cerebellum mirrors that in the cerebral cortex

The strong contribution of the visual system in mode 1 and 2 is particularly interesting (Fig.1,2 and S1,2 Fig A, B). Although the primary visual representation remained traditionally undetected and was conceptually denied in earlier brain-imaging research on the human cerebellum based on resting-state functional connectivity (Buckner et al., 2011), recently, van Es et al. (2019) identified three distinct visual spatial representations in the cerebellum (oculomotor vermis, lobule VIIb, lobule VIIIb). In resting-state fMRI data from HCP, Guell et al. (2020) identified a representation of the “visual functional territory” that was separated from other functional territories of the human dentate nucleus. Xue et al. (2021) recently confirmed the early retinotopic visual cortex representation in detailed cerebellar parcellation in three human subjects. Our present results add evidence that speak to a visual cortical representation in the cerebellum, from the novel angle of structural covariation at the population level.

Further, King et al. (2019) also treated lobule VI vermis as a region related to complex saccades that require attentional control. Indeed, there is some evidence that the DAN and the visual network representation can be closely associated in the cerebellum (Brissenden et al., 2016; King et al., 2023). King et al. (2023) showed that the oculomotor vermis of the cerebellum is predicted best by neural activity responses of regions in DAN as well as the extrastriate visual cortex. Brissenden et al. (2018) also showed visual field representation in lobule VIIb/VIIIa that mirrors a portion of IPS, which is considered part of the DAN. Moreover, the DAN has a close relationship with sensorimotor regions and serves a key role in the visuospatial perceptual attention (Buschman & Kastner, 2015; Corbetta & Shulman, 2002; Dixon et al., 2018; Ptak, 2012).

We note that the DAN was virtually absent from mode 1 (Funct7net, Struct7net and Funct100region). DAN also had opposite ties to other higher-order cognitive networks in mode 2 (Funct7net, Struct7net and Funct100region). These higher-order cognitive systems included the default mode network (DMN), ventral salience network (VSN), limbic network and executive control network (ECN). While there is abundant evidence that DAN exhibited a different pattern compared to other higher-order cognitive systems in the cerebral cortex, our present findings suggest that DAN might also be anchored in a unique way in the cerebellum compared to the other higher-order cognitive systems. Cortical DAN and default mode network (DMN) typically show anticorrelated relationships, which has been widely recognized as a general feature of human brain organization (Andrews-Hanna et al., 2014; Fox et al., 2005; Spreng, 2012). Our findings suggest that this antagonistic relationship may also manifest in the human cerebellum.

### Ipsilateral cerebellum-cortex correspondence in humans can be important

Whereas cerebellum-cortex communications are usually portrayed as *contralateral*, our findings in mode 3 (Funct 100region) identified under-appreciated *ipsilateral* cerebrocerebellar interactions. The feedforward pathways linking cerebral cortex to cerebellum are conveyed in cortico-ponto-cerebellar (CPC) projections that originate from the frontal, temporal, parietal and occipital lobes of the cerebral cortex, and terminate on nuclei of the ipsilateral basis pontis. Pontocerebellar axons then traverse the pons to enter the contralateral middle cerebellar peduncle and course to the contralateral cerebellum. Notably, up to 30% of these axons cross again in the cerebellar white matter to terminate on the cerebellum on the same side as the cerebral cortical areas of origin (Na et al., 2019; Rosina & Provini, 1984). The feedback pathways from cerebellum to cerebral cortex are conveyed in the dentato-rubro-thalamo-cortical tracts (DRTT or DRTC) that originate in the deep cerebellar nuclei (dentate nucleus, but also the fastigial and the interposed – globose and emboliform nuclei), course in the superior cerebellar peduncle, decussate in the brachium conjunctivum, pass through the contralateral red nucleus to which they provides passing collaterals, and then ascend to terminate in a discrete set of nuclei in the thalamus. The thalamocortical projections close the feedback circuit (Voogd, 2004). Anatomical tract tracing studies in animals show that the major connections between the cerebellum and cerebral cortex are contralateral (Kelly & Strick, 2003; Schmahmann & Pandya, 1989b, 1991b, 1997b; Wiesendanger & Wiesendanger, 1985), although there are some ipsilateral or non-decussating DRTT (nd-DRTT) pathways in monkeys (Chan-Palay, 1977; Wiesendanger & Wiesendanger, 1985), cats (Flood & Jansen, 1966), and ∼ 20% ipsilateral DRTT and pontocerebellar pathways in rats (Aumann & Horne, 1996; Cicirata et al., 2005; Serapide et al., 2002). As argued by Doron et al. (2010) and others, assumptions about anatomical connections in the human brain inferred from the tract-tracing studies performed in monkeys should be considered with care, as the association cortices are more developed and may project more to the cerebellum in humans than in animals (Axer & Keyserlingk, 2000; Ramnani et al., 2006).

Ipsilateral pathways are evident also in studies of the human cerebellum. In the feedforward system, DTI studies suggest that there are ipsilateral CPC tracts, traveling from the temporal lobe, occipital lobe (Karavasilis et al., 2019; Keser et al., 2015; Palesi et al., 2017; Sokolov et al., 2014) and parietal lobe (Karavasilis et al., 2019; Keser et al., 2015) to the cerebellum. Microsurgical human post-mortem brain microdissections of the feedback projection reveal an nd-DRTT pathway ((Meola et al., 2016; Tacyildiz et al., 2021), and DTI fiber tract reconstructions in the living brain identified an nd-DRTT pathway accounting for one-fifth of the overall DRTT (Karavasilis et al., 2019; Keser et al., 2015; Meola et al., 2016; Petersen et al., 2018). Our own data would be in line with these ipsilateral cerebellum-cortex correspondences (Fig. 3). Despite DTI’s ability to reconstruct the major contralateral feedforward and feedback pathways (Kamali et al., 2010; Karavasilis et al., 2019; Keser et al., 2015; Meola et al., 2016; Palesi et al., 2017; Petersen et al., 2018), it is inherently limited in resolving crossing and kissing fibres. Thus, the interpretation of DTI results can be challenging (Habas & Cabanis, 2007) and this may impact statements on hemispheric asymmetry. Support for the ipsilateral connectivity in humans, however, comes from brain stimulation studies. Repetitive transcranial magnetic stimulation (rTMS) and transcranial direct current stimulation (tDCS) on one side of the cerebellum produce strong bilateral effects on fMRI or PET studies of motor tasks (Miall & Christensen, 2004) and verbal working memory retrieval tasks (Macher et al., 2014) as well as brain-wide cortical metabolism (Cho et al., 2012). Further, in resting-state functional connectivity studies in humans, Igelstrom et al. (2017) noted functional connectivity between seed regions on one side of the temporoparietal junction (TPJ) and bilateral areas in Lobules V, VI, Crus I/II, Lobule VIIIa/b, and IX. The emphasis was on the contralateral side, but the ipsilateral functional connectivity between each seed region and cerebellum was prominent.

Our ipsilaterally patterned modes document ipsilateral correspondence underlying the population covariation between the cerebellum and the cortex on a large scale. Our structural covariation approach is applicable at the whole-brain level and unchangeable because it is applied purely to structural data and independent of the experiments or subject outliers. This provides a degree of confidence in the validity of the observations, the significance of which remains to be determined in future research.

### PheWAS revealed distinct cerebellar and cortical phenotypic profiles for each mode

Our PheWAS has screened a diverse palette of phenotypic indicators to delineate their associations with our derived cortical and cerebellar brain patterns. Within each mode, numerous phenotypes associated with either cortical or cerebellar latent variables have also demonstrated significant brain-behavior links.

For the cerebellar mode 1 (Funct7net and Funct100region; Fig. 6), we found the cognitive phenotypes measuring psychomotor speed, physical activity, fat-free mass, hand grip strength, blood cell metrics and cardiovascular measurement to be particularly relevant. The cognitive tests indexing psychomotor processing speed (e.g., reaction time, DSST and TMT-A) and hand grip strength showed robust and significant association with the cerebellar mode 1. The strong association between these phenotypes was reported in another UKBB study (Jiang et al., 2022). Multiple studies across different populations indicated physical exercise is positively correlated with cerebellar grey matter volume (Ben-Soussan et al., 2015; Li et al., 2022), gray matter mean diffusivity (Callow et al., 2021) and network connectivity (Won et al., 2021) coupled with improved cognitive performance. In rats, exercise induces angiogenesis in the cerebellum, which resonates with our strong association between blood cells (red blood cells, white blood cells, reticulocytes and IGF-1) and cerebellum (Black et al., 1990; Isaacs et al., 1992; Lopez-Lopez et al., 2004). However, gamma-glutamyl transferase (GGT) and urate’s association in our findings with the cerebellum was unclear. The cerebellum is thought to assist in regulating autonomic functions such as heart rate, respiration and sleep-wake cycle through hindbrain projections from parabrachial nuclei (PBN) (Chen et al., 2023; Hashimoto et al., 2018; Martner, 1975; Moruzzi, 1947; Nisimaru, 2004; Nisimaru & Katayama, 1995; Nisimaru et al., 2013; Rasheed et al., 1970; Reis et al., 1973; Zanchetti & Zoccolini, 1954) Thus, diastolic blood pressure (DBP) and ventricular rate could indeed be associated with individual differences in cerebellar morphology. In cerebellar mode 2 (Funct7net), we also identified sleep-related phenotypes and snoring, which could be regulated through this pathway as well (Canto et al., 2017; DelRosso & Hoque, 2014; Hashimoto et al., 2018; Pedroso et al., 2011). Previous studies observed a relationship between height and cerebellum volume in adults (Hara et al., 2016; Hutchinson et al., 2003; Park et al., 2009; Taki et al., 2012). Similarly, BMI has strong associations with gray matter volume in middle-aged and older adults, including the cerebellum (Kurth et al., 2013; Pannacciulli et al., 2006; Taki et al., 2008; Walther et al., 2010). In particular, Weise et al. (2013) found that although both fat mass and fat-free mass (FFM) contribute to BMI, they have differential impacts on brain structure, including a negative association between cerebellar GMV and fat mass, but not FFM. Since FFM consists mainly of skeletal muscle, bones and parenchymal organs which are metabolically active, this echoes a close link of exercise with the cerebellum, in line with our analysis.

For the cortical mode 1 (Funct7net), the various cognitive phenotypes related to abstract reasoning, executive functions, focused visual attention and motor control confirm the known functions of the cerebral cortex, and highlight other phenotypes including time spent watching TV, fat mass, blood indicators of diabetes, immunoreactions, and cardiovascular disease. For example, TV consumption played an important role in both mode 1 and mode 2 (Funct7net). TV consumption is both a sedentary and passive-receptive behavior. For mode 1, a wide range of studies have shown that excessive TV watching was negatively correlated with gray matter volume in multiple cortical regions and cognitive functions in both the young and elderly (Dougherty et al., 2022; Takeuchi & Kawashima, 2023; Takeuchi et al., 2015) but positively correlated with risks of type-2 diabetes, cardiovascular diseases and increased BMI (Grøntved & Hu, 2011; Patterson et al., 2020), despite physical activity (Chomistek et al., 2013; Ekelund et al., 2016). The blood traits are strongly confirmatory of these results, as we saw HbA1c which measures blood sugar level and is a marker for diabetes. We here also observed LDL and ApoB links, which were highly associated with sedentary time (Crichton & Alkerwi, 2015) and are important predictors of cardiovascular diseases (Contois et al., 2023; Sniderman et al., 2022; Sniderman et al., 2011). In addition, elevated basophil count was reported in some diabetic patients (Shah et al., 2021) and coronary artery disease patients (Pizzolo et al., 2021). The effects of basophil might be related to adiposity-associated inflammatory factors (Zhang et al., 2021). Finally, Systolic blood pressure and carotid intima-media thickness show significant association with the cerebral cortex, which is consistently observed across different types of studies and populations (Ferreira et al., 2016; Juonala et al., 2010; Liu et al., 2017; Zhang et al., 2019). Interestingly, high lipoprotein levels and diabetes as seen in blood traits for the cortex, obesity and a sedentary lifestyle as seen in physical and lifestyle phenotypes for the cortex, all contribute to the thickening of carotid intima-media.

For the cortical mode 2 (Funct7net; S5) which separates the visual attention system from other higher-order cognitive systems (cf. above), we identified watching TV as the leading phenotype associated with both the cortex and the cerebellum – potentially another piece of evidence for complex behaviors being related to the cerebellum. Apart from being sedentary, watching TV’s more receptive nature differentiates it from behaviors such as reading or gaming. Many observed that watching TV activated key nodes in DMN and suppressed DAN (Anderson & Davidson, 2019; Raichle, 1998) which is coherent with our interpretation of mode 2 (Funct7net) that this mode reflects the anticorrelation between DAN and DMN. In fact, attentional deficits were often observed in children and adolescents with too much TV exposure (Christakis et al., 2004; Johnson et al., 2007; Landhuis et al., 2007). At the same time, one receives abundant visual and narrative stimuli while watching TV, which is directly associated with the visual system and with semantic processing in association cortex (Yang et al., 2023). Thus, cerebellar structural brain variation can be expected to be jointly reflected in visual and association cortices, an idea which receives support from a cross-sectional study on TV viewing and brain structure (Takeuchi et al., 2015).

In contrast to mode 1, mode 2’s significant cognitive phenotypes mostly relate to fluid intelligence tests and prospective memory. Both faculties closely implicate executive functions and working memory, which are supported by the visual-attention system (Burgess et al., 2001; Cochrane et al., 2019; Engle et al., 1999; Kliegel et al., 2002). Moreover, the fluid intelligence tests were characterized by arithmetical tasks, which demand effortful top-down visual attentional control (Wong & Liu, 2020).

In both mode 1 and 2 (Funct7net), we noticed smartphone usage was most associated with cerebral cortex variation. Excessive smartphone usage is related to altered structures in frontal, temporal, insular and anterior cingulate regions and reduced neural activity responses when viewing emotional stimulus in fronto-cingulate regions (Chun et al., 2017; Horvath et al., 2020; Tymofiyeva et al., 2020; Wang et al., 2016). These regions were implicated in cerebellar-cortical population modes as part of the higher cognitive systems that stands in some antagonistic contrast with lower sensorimotor system.

In conclusion, our first two modes are consistent with the DMN-visuomotor divergence of neural systems, as well as the classical double motor representation and the recently proposed triple nonmotor representation in the cerebellum. Our results support the anticorrelation between visual-attention and other higher order cognitive systems in the cerebellum, like in the cortex. Our third mode indicates a greater proportion of ipsilateral mechanisms between the cerebellum and cortex that may be overshadowed by the more abundant contralateral pathways. The distinct phenotype profiles for each mode and their latent variables revealed unique brain phenomena - behavior links that weighed differently in the cerebellum and the cortex. These findings greatly contribute to our understanding of the intricate interplay among the cortex, cerebellum and behaviors. Currently, the analysis was done on a cross-sectional cohort and only involved the cerebral cortex. In the future, we plan to investigate the longitudinal effect of cerebellar-cortical structural covariation and potentially study the subcortico-cerebellar interactions for a more complete picture.

## Acknowledgements

BTTY is supported by the NUS Yong Loo Lin School of Medicine (NUHSRO/2020/124/TMR/LOA), the Singapore National Medical Research Council (NMRC) LCG (OFLCG19May-0035), NMRC CTG-IIT (CTGIIT23jan-0001), NMRC STaR (STaR20nov-0003), Singapore Ministry of Health (MOH) Centre Grant (CG21APR1009), the Temasek Foundation (TF2223-IMH-01), and the United States National Institutes of Health (R01MH120080 & R01MH133334). Any opinions, findings and conclusions or recommendations expressed in this material are those of the authors and do not reflect the views of the Singapore NMRC, MOH or Temasek Foundation.

