## Supplemental Material for "Intrinsic structural covariation links cerebellum subregions to the cerebral cortex"

A
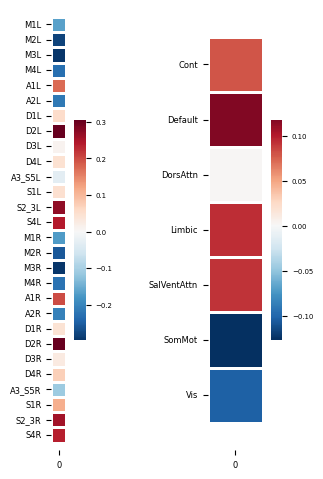
 B
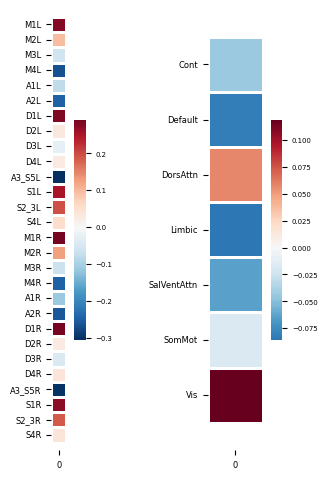

**Supplementary Figure 1: Funct7net Brain weights for Mode 1 and Mode 2.** Within each pair of weights, cerebellar weights were shown on the left and cortical weights were shown on the right. This analysis was based on ~38,527 participants in the UK Biobank dataset. **A)** For Mode 1, bilateral eye (M1L/R), mouth (M2L/R), upper limbs (M3L/R), lower limbs (M4L/R) and action observation (A2L/R) regions in the cerebellum showed the same direction of structural covariation with the cortical somatomotor (SomMot) and visual (Vis) networks. In contrast, bilateral executive (D2L/R), social and language (S2_3L/R, S4L/R) and complex action (A1L/R) regions in the cerebellum showed the same direction of structural covariation with cortical Executive Control (Cont), Default, Limbic and Ventral Attention (SalVentAttn) Network. Dorsal Attention Network (DorsAttn) had nearly zero weights. **B)** For Mode 2, bilateral eye (M1L/R), executive (D1L/R), social and language (S1L/R, S2_3L/R) and mouth (M2L/R) regions in the cerebellum showed the same direction of structural covariation with the cortical Dorsal Attention Network (DorsAttn) and visual (Vis) networks. In contrast, bilateral action observation + introspection (A3_S5L/R), action observation (A2L/R) and lower limbs (M4L/R) showed the same direction of structural covariation with the cortical Default, Limbic, Ventral Attention (SalVentAttn) and Executive Control (Cont) Networks. The hot and cold colors do not represent directionality but simply differentiate two opposite directions of structural covariations. L= Left cerebellum hemisphere, R= Right cerebellum hemisphere.

A
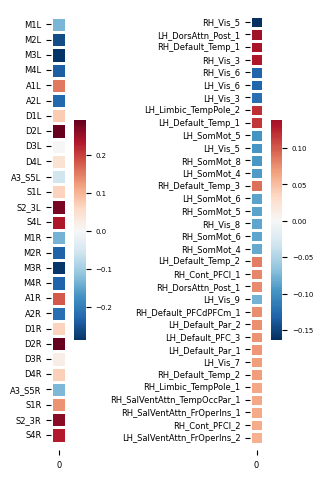
B
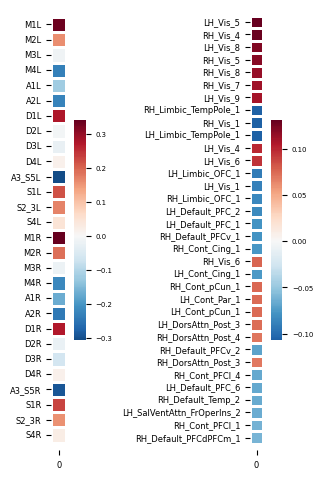

C
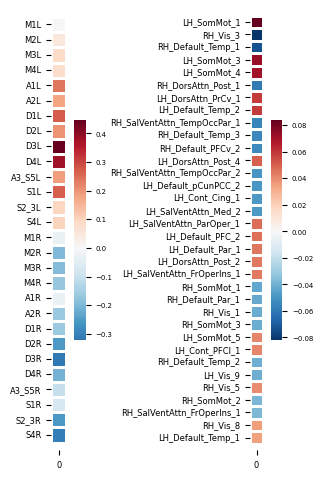

**Supplementary Figure 2: Funct100region Brain weights for Mode 1, Mode 2 and Mode 3.** Within each pair of weights, cerebellar weights were shown on the left and cortical weights were shown on the right. This analysis was based on ~38,527 participants in the UK Biobank dataset. The cerebellar weights were highly similar to the cerebellar weights in Funct7net Mode 1 and Mode 2. The cortical and cerebellar weights for Mode 3 show the ipsilaterally same direction of structural covariation. For the cortical weights, we only showed the top 34 regions out of 100 regions with the highest absolute weights. **A)** In Mode 1, the cortical regions sharing the same direction of structural variation with higher-order cognitive systems are distributed in LH_DorsAttn_Post_1 (left visual word form area / fusiform gyrus), RH_Default_Temp_1 (right middle-inferior temporal gyrus), RH_Vis_3 (right visual word area/ fusiform gyrus), LH_Limbic_TempPole_2 (left inferior temporal lobe), LH_Default_Temp_1 (left anterior temporal pole), RH_Default_Temp_3(right Superior temporal sulcus / auditory cortex), LH_Default_Temp_2 (left superior temporal sulcus / auditory cortex) among other regions. In contrast, the cortical regions sharing the same direction of structural variation with lower visual-motor systems are distributed in RH_Vis_5 (right primary visual cortex), RH_Vis_6 (right retrosplenial/parahippocampal cortex), LH_Vis_6 (left retrosplenial cortex), LH_Vis_3 (left lingual gyrus / fusiform gyrus), LH_SomMot_5(left primary motor cortex for hand), LH_Vis_5 (left primary visual cortex), RH_SomMot_8(right primary/supplementary motor cortex) LH_SomMot_4 (left premotor cortex for speech production) among other regions. **B)** In Mode 2, the cortical regions sharing the same direction of structural variation with visual-attention system are distributed in L/RH_Vis_4 (middle-inferior occipital gyrus) /5 (primary visual cortex) /6 (PreCuneus/PCC) /8 (middle occipital gyrus), RH_Vis_7 (right extrastriate cortex), LH_Vis_9 (cuneus), L/RH_Cont_pCun_1 (precuneus), R/LH_DorsAttn_Post_3/LH_Cont_Par_1/ RH_DorsAttn_Post_4 (intraparietal sulcus) among other regions. In contrast, the cortical regions sharing the same direction of structural variation with other higher-order cognitive systems are distributed in L/RH_Limbic_TempPole_1 (anterior temporal pole), L/RH_Vis_1 (parahippocampal cortex), L/RH_Limbic_OFC_1(orbitofrontal cortex), LH_Default_PFC_1(anterior insula)/2(inferior frontal gyrus), RH_Default_PFCv_1(right orbitofrontal/ventrolateral cortex), L/RH_Cont_Cing_1(posterior cingulate cortex), RH_Default_PFCv_2(right inferior frontal cortex), RH_Cont_PFCl_4(right dorsolateral prefrontal cortex) among other regions. **C)** In Mode 3, left complex action (A1L), action observation (A2L), executive (D1/2/3/4L), action observation + introspection (A3_S5L) and social and language (S1L) displayed the strongest weights on the left side of the cerebellum, sharing the same direction of structural covariation with multiple regions in the left side of the cortex, including LH_SomMot_1(primary auditory cortex, superior temporal gyrus, heschl gyrus, planum temporale), LH_SomMot_3(somatosensory cortex), LH_SomMot_4(premotor cortex), LH_DorsAttn_PrCv_1(premotor cortex), LH_Default_Temp_2(superior temporal sulcus / auditory cortex), LH_DorsAttn_Post_4(anterior intraparietal sulcus), LH_SalVentAttn_ParOper_1(supramarginal gyrus), LH_Default_PFC_2(inferior frontal gyrus) among other regions. In contrast, right mouth(M2R), upper limbs (M3R), lower limbs (M4R), action observation (A2R), executive (D1/2/3/4R), social and language (S2_3R, S_4R) displayed the strongest weights on the right side of the cerebellum, sharing the same direction of structural covariation with multiple regions in the right side of the cortex, including RH_Vis_3(visual word area/ fusiform gyrus), RH_Default_Temp_1(right middle-inferior temporal gyrus), RH_DorsAttn_Post_1(temporoparietal junction), RH_SalVentAttn_TempOccPar_1(posterior superior temporal sulcus/planum temporale), RH_Default_Temp_3 (right Superior temporal sulcus / auditory cortex), RH_Default_PFCv_2(right inferior frontal cortex), RH_SalVentAttn_TempOccPar_2(primary somatosensory cortex). Notable exceptions to this ipsilaterally same direction of structural covariation are bilateral precuneus/posterior cingulate cortices. The hot and cold colors do not represent directionality but simply differentiate two opposite directions of structural covariations. L= Left cerebellum hemisphere, R= Right cerebellum hemisphere. LH=Left cortex, RH= Right cortex. The cortical regions are functionally decoded from Neurosynth.

A
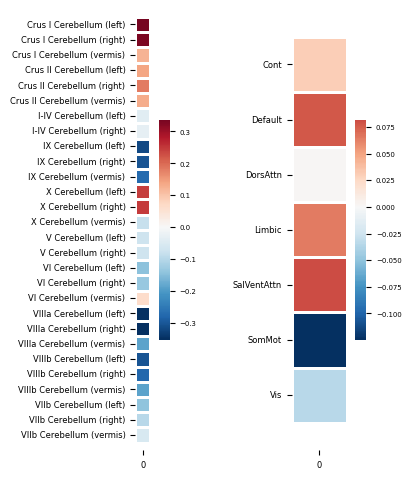
 B
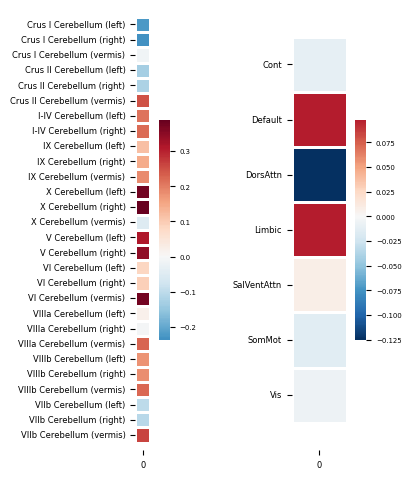

C
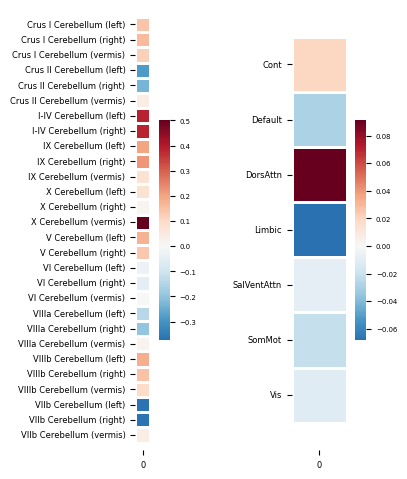
D
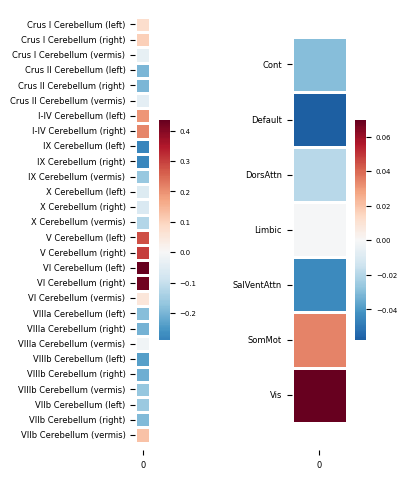

**Supplementary Figure 3: Struct7net** **Brain weights for (A) Mode 1, (B) Mode 2, (C) Mode 3, and (D) Mode 4.** Within each pair of weights, cerebellar weights were shown on the left and cortical weights were shown on the right. This analysis was based on ~38,527 participants in the UK Biobank dataset. Mode 1 and Mode 4 exhibited highly similar cortical weights to Funct7net Mode 1’s cortical weights. Mode 2 captured the anticorrelation between DMN and DAN shown in Funct7net and Funct100region Mode 2. For a description of each mode, see the main text Fig 3. The hot and cold colors do not represent directionality but simply differentiate two opposite directions of structural covariations.

A
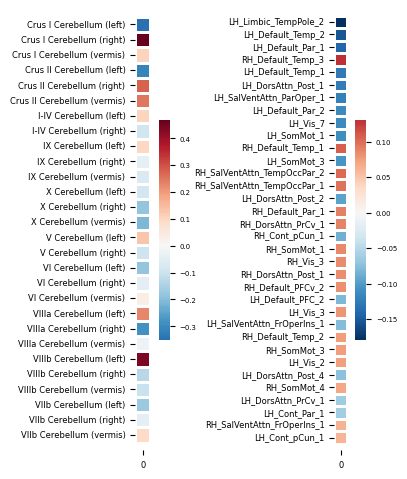
B
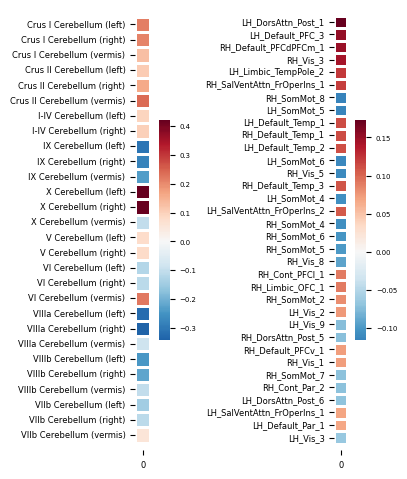

C
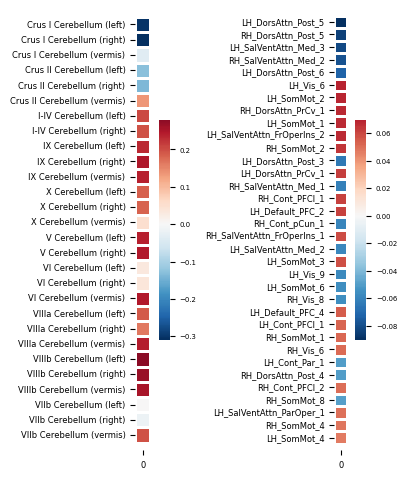

**Supplementary Figure 4: Struct100region** **Brain weights for (A) Mode 2, (B) Mode 3 and (C) Mode 5.** Within each pair of weights, cerebellar weights were shown on the left and cortical weights were shown on the right. This analysis was based on ~38,527 participants in the UK Biobank dataset. Mode 2 also exhibited an ipsilaterally same direction of structural covariation between the most significant regions in the cerebellum and the cortex. Mode 3 exhibited the higher-order cognitive system vs lower sensorimotor system interplay seen in Funct7net/Funct100region Mode 1, and Struct7net Mode 1 and Mode 4. Mode 5 exhibited the DMN-DAN anticorrelation seen in Funct7net/Funct100region/Struct7net Mode 2. The hot and cold colors do not represent directionality but simply differentiate two opposite directions of structural covariations. LH=Left cortex, RH= Right cortex.

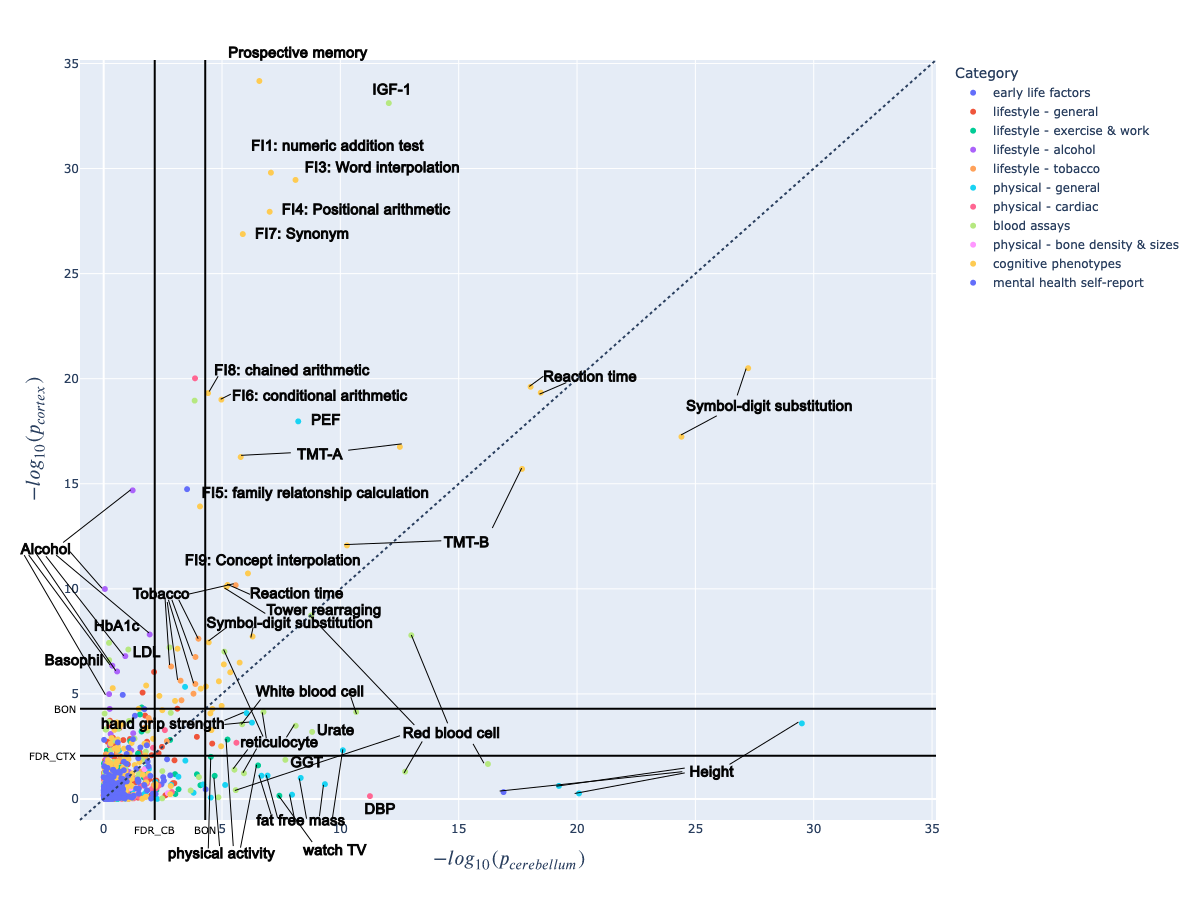

**Supplementary Figure 5: Funct100region Mode 1’s phenotype array was mostly similar to Funct7net Mode 1. IGF-1’s association with the cortex increased significantly, while several other blood essays decreased. Expiratory measurements and tobacco also showed increased association with the cortex than the cerebellum. SBP and CIMT became insignificant.** The x-axis showed the -log_10(p-value) of the cerebellum while the y-axis showed the -log_10(p-value) of the cortex. The Bonferroni correction threshold (BON) and False Discovery Rate are plotted for the cortex and the cerebellum on the respective axis. Phenotypes near the upper-left part of the figure are more strongly associated with the cortex while those near the lower-right corner are more strongly associated with the cerebellum. The phenotypes near the diagonal are relevant for both the cortex and the cerebellum. The phenotypes at the lower-left corner bordered by the two BON threshold lines are considered insignificant.

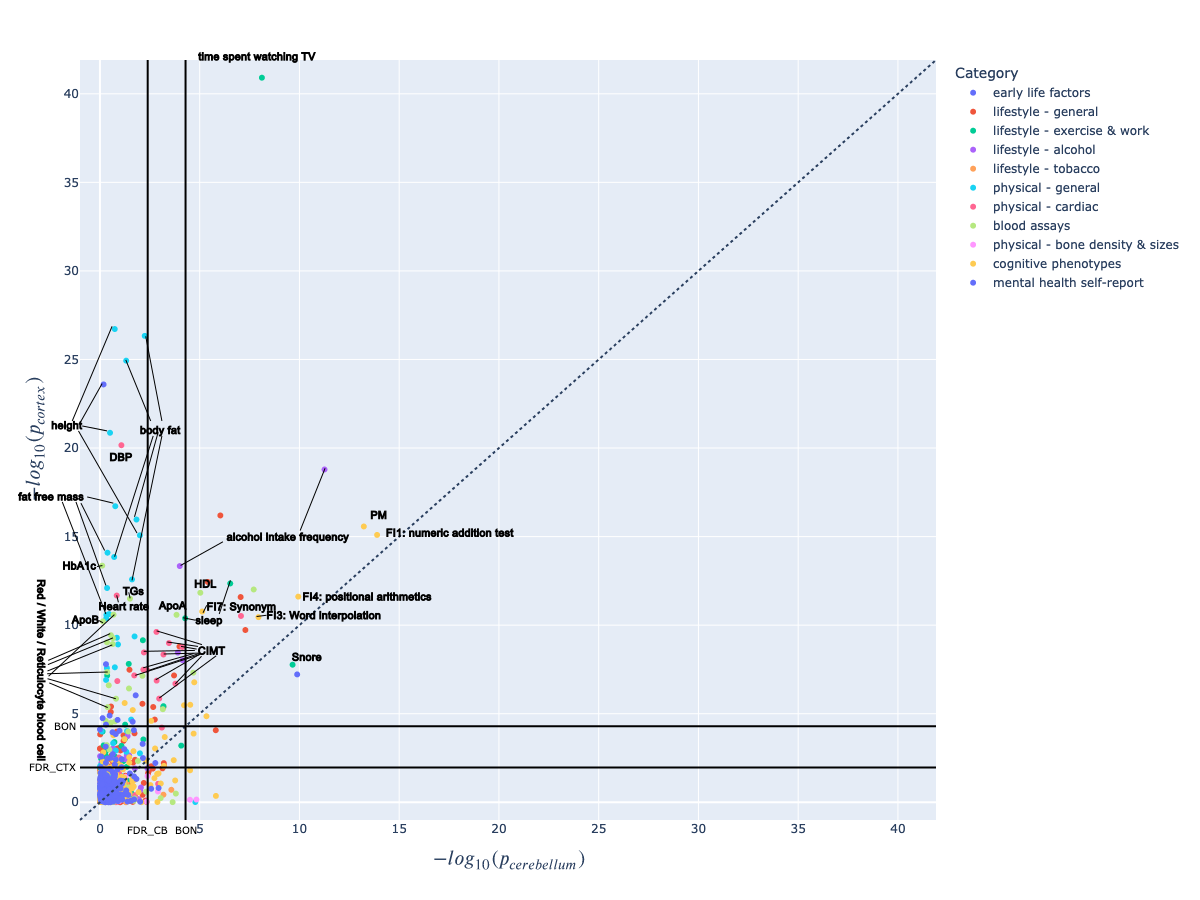

**Supplementary Figure 6: Funct100region Mode 2’s phenotype array was mostly similar to Funct7net Mode 2.** The x-axis showed the -log_10(p-value) of the cerebellum while the y-axis showed the -log_10(p-value) of the cortex. The Bonferroni correction threshold (BON) and False Discovery Rate are plotted for the cortex and the cerebellum on the respective axis. Phenotypes near the upper-left part of the figure are more strongly associated with the cortex while those near the lower-right corner are more strongly associated with the cerebellum. The phenotypes near the diagonal are relevant for both the cortex and the cerebellum. The phenotypes at the lower-left corner bordered by the two BON threshold lines are considered insignificant.

**A**
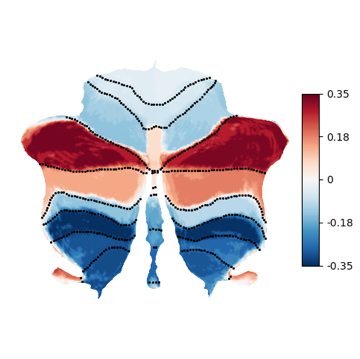

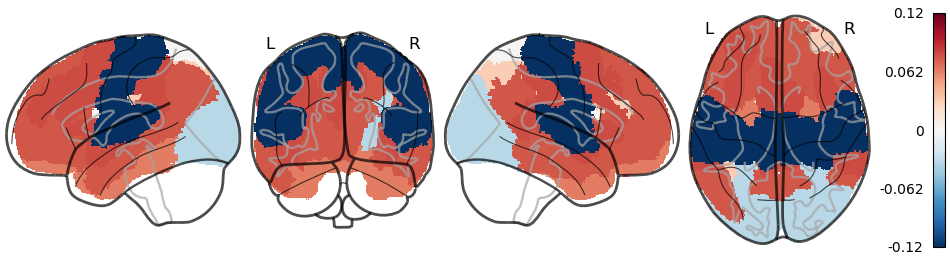

**B**
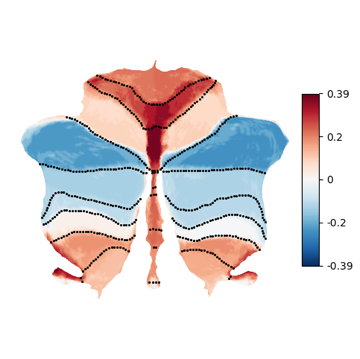

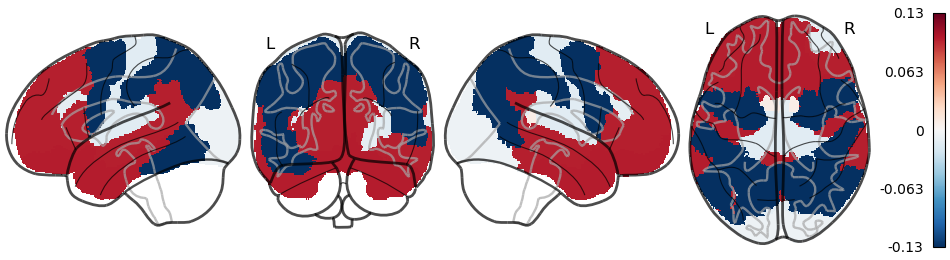

**C**
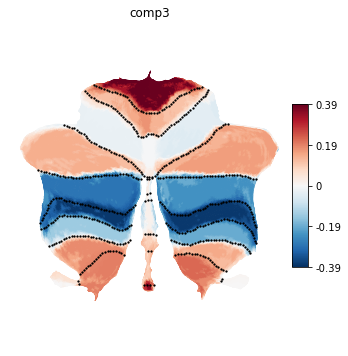

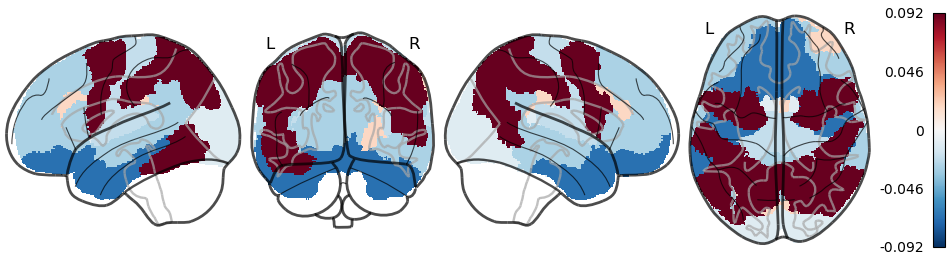

**D**
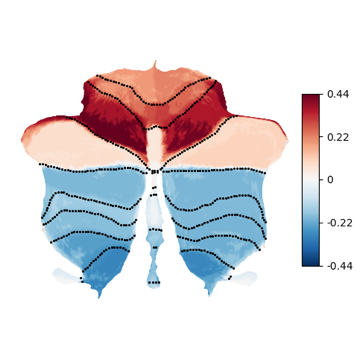

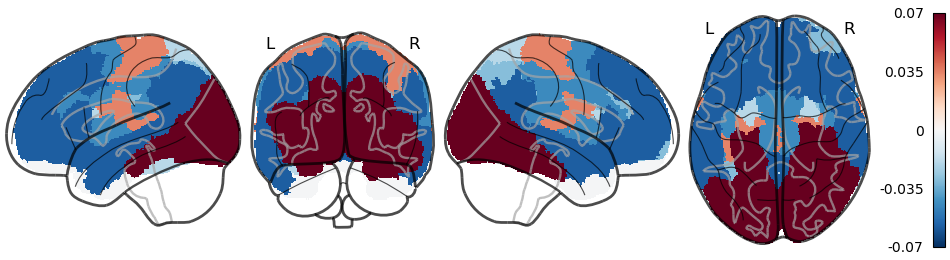

**Supplementary Figure 7. Three complementary modes of structural covariation between 28 structural cerebellar regions and 7 cortical networks (Struct7net).** Mode 1 and Mode 4 in cerebellar structural parcellation with cortical 7 networks averaged ('Struct 7net') recapitulated the contrast seen in 'Funct 7net' Mode 1 (Fig.1A) with highly correlated cortical weights (Mode 1: r = 0.94, p-value < 10^-47; Mode 4: r = -0.92, p-value < 10^-42). Mode 2 in 'Struct 7net' revealed a strong anti-correlation between the dorsal attention network (DAN) and the default mode network (DMN) / limbic network, supporting the notion of an antagonist relationship anchored in the cerebellum. **(A)** Mode 1 had an explained variance of 0.20 and had the most coherent cortical weights with 'Funct 7net' Mode 1. We observed one end of structural variation weights in crus I/II and bilateral lobule X. The other end of structural variation weights was seen in lobules VIIIa/b, lobules IX, and the anterior lobe to a lesser extent. **(B)** Mode 2 had an explained variance of 0.19. One end of structural variation weights was observed in the anterior lobe, vermis across the posterior lobe, bilateral lobules VIIIb, and lobule X in the cerebellum and DMN and limbic networks in the cortex. The other end of structural variation was observed in bilateral lobules VII and DAN. **(C)** Mode 3 had an explained variance of 0.13. one end of structural variation weights was primarily seen in lobule X vermis, anterior lobe, bilateral lobules VIIIb and IX and Crus I, while the other end were in bilateral lobules VIIb, Crus II and lobules VIIIa. **(D)** Mode 4 had an explained variance of 0.12 and also resembles 'Funct 7net' Mode 1’s cortical weights, except for the limbic network. One end of structural variation weights was primarily seen in the anterior lobe, with some involvement of crus I. The other end of structural variation weights was mainly seen in the posterior lobe starting from crus II, excluding most of the cerebellar vermis or lobule X.

**A**
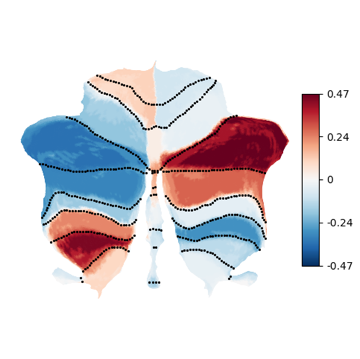

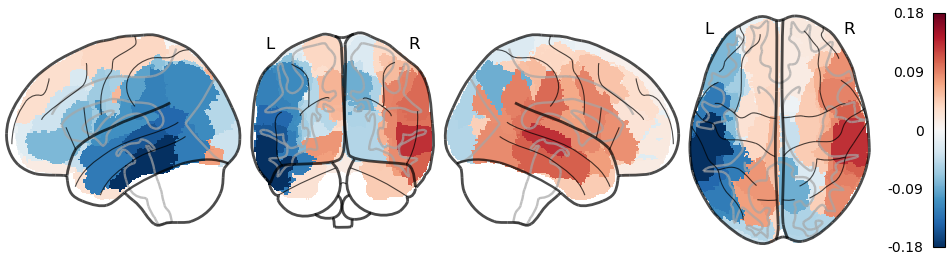

**B**
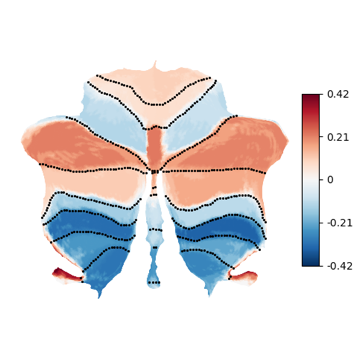

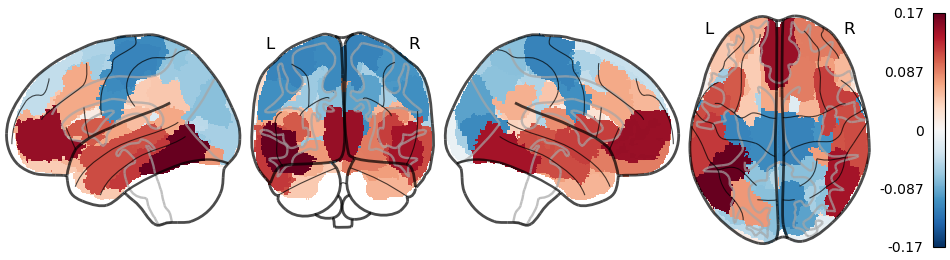

**C**
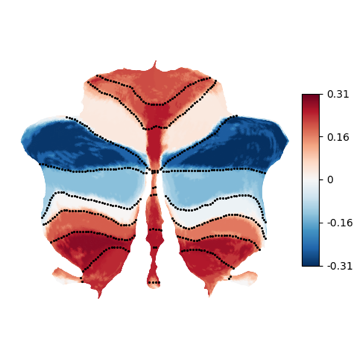

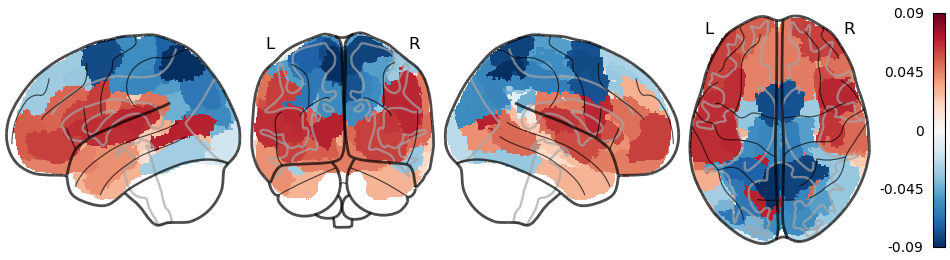

**Supplementary Figure 8. Mode 2, Mode 3 and Mode 5 of structural covariation between 28 structural cerebellar regions and 100 cortical regions (Struct100region) reconstructed the ipsilateral pattern, the high vs. low pattern and the DAN-DMN/Limbic anticorrelation respectively.** (A) Mode 2 in ‘Struct 100region’ has an ipsilatetal pattern and can be compared to ‘Funct 100region’ Mode 3 with extremely similar cortical weights. **(B)** Mode 3 in cerebellar structural parcellation with cortical 100 region parcellation (‘Struct 100region’) corresponded to ‘Funct 100region’ Mode 1 (Fig.2A) and ‘Struct 7net’ Mode 1 (Fig.3A), with highly correlated cortical weights (r=0.85, p-value < 10^-27) and cerebellar weights (r=0.93, p-values < 10^-11) respectively. It had an explained variance of 0.27. **(C)** Mode 5 in ‘Struct 100region’ corresponded to ‘Funct 100region’ Mode 2 (Fig.2B) and ‘Struct 7net’ Mode 2 (Fig.3B) with significantly different cortical weights but highly correlated cerebellar weights (r=0.84, p-values < 10^-7). DAN hubs and DMN / Limbic regions dominated two ends of the structural variation.

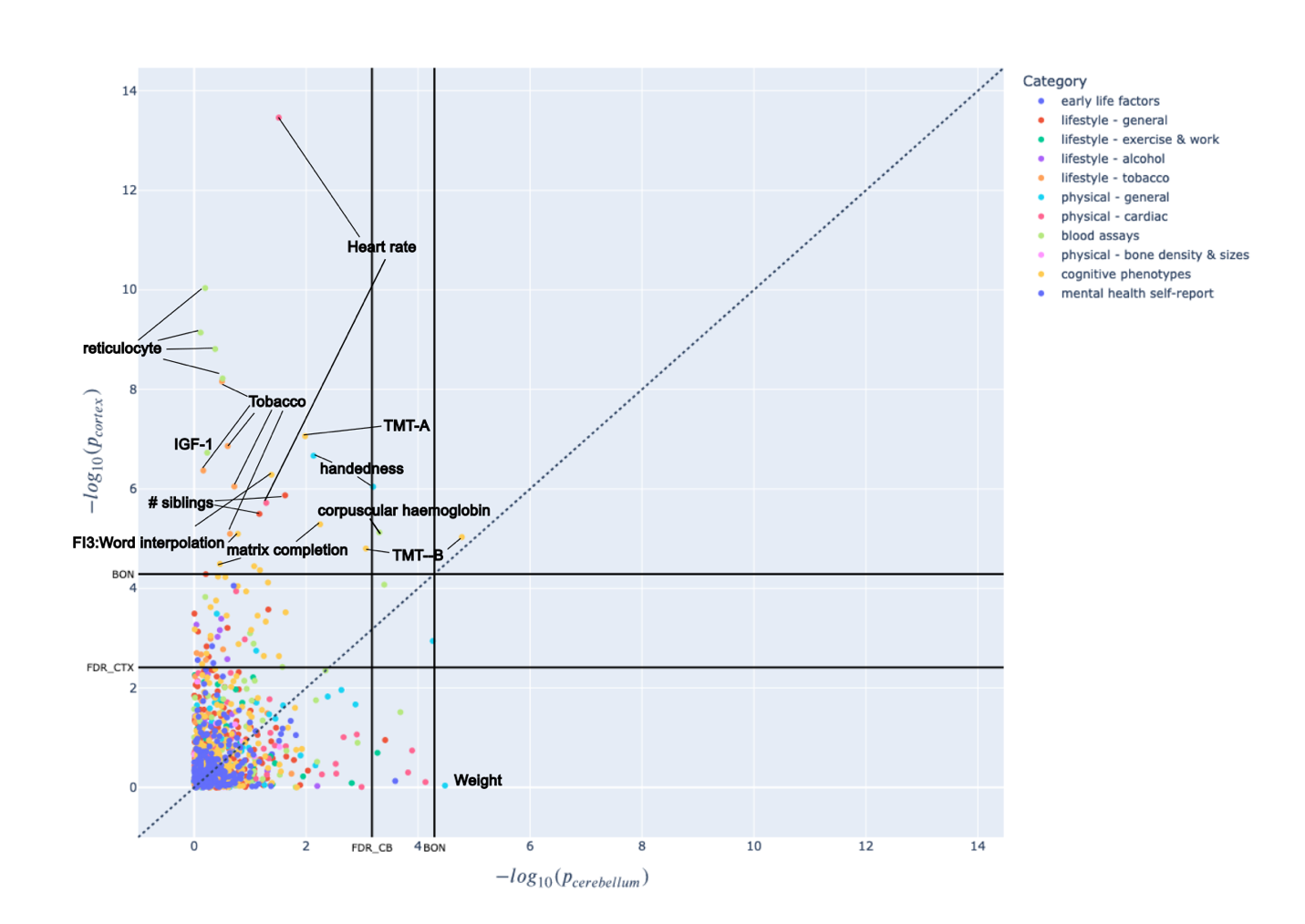

**Supplementary Figure 9: Funct100region Mode 3’s phenotype array.**

| label | side | function |
| --- | --- | --- |
| M1L | left | eye |
| A1L | left | mouth |
| M3L | left | hands |
| M4L | left | lower |
| M2L | left | complex action |
| A2L | left | action observation |
| D1L | left | executive |
| D2L | left | executive |
| D3L | left | executive |
| D4L | left | executive |
| A3_S5L | left | action observation + introspection |
| S1L | left | social and language |
| S2_3L | left | social and language |
| S4L | left | social and language |
| M1R | right | eye |
| A1R | right | mouth |
| M3R | right | hands |
| M4R | right | lower |
| M2R | right | complex action |
| A2R | right | action observation |
| D1R | right | executive |
| D2R | right | executive |
| D3R | right | executive |
| D4R | right | executive |
| A3_S5R | right | action observation + introspection |
| S1R | right | social and language |
| S2_3R | right | social and language |
| S4R | right | social and language |

**Supplementary Table 1: Functionally derived cerebellar atlas region labels and functions.**
